# A Refined View of Airway Microbiome in Chronic Obstructive Pulmonary Disease at Species and Strain-levels

**DOI:** 10.1101/2020.01.18.908624

**Authors:** Zhang Wang, Haiyue Liu, Fengyan Wang, Yuqiong Yang, Xiaojuan Wang, Boxuan Chen, Martin R. Stampfli, Hongwei Zhou, Wensheng Shu, Christopher E. Brightling, Zhenyu Liang, Rongchang Chen

## Abstract

Little is known about the species and strain-level diversity of the airway microbiome, and its implication in chronic obstructive pulmonary disease (COPD).

Here we report the first comprehensive analysis of the COPD airway microbiome at species and strain-levels. The full-length 16S rRNA gene was sequenced from sputum in 98 stable COPD patients and 27 age-matched healthy controls, using the ‘third-generation’ Pacific Biosciences sequencing platform.

Individual species within the same genus exhibited reciprocal relationships with COPD and disease severity. Species dominant in health can be taken over by another species within the same genus in GOLD IV patients. Such turnover was also related to enhanced symptoms and exacerbation frequency. *Ralstonia mannitolilytica*, an opportunistic pathogen, was significantly increased in COPD frequent exacerbators. There were inflammatory phenotype-specific associations of microbiome at the species-level. One group of four pathogens including *Haemophilus influenzae and Moraxella catarrhalis*, were specifically associated with sputum mediators for neutrophilic inflammation. Another group of seven species, including *Tropheryma whipplei*, showed specific associations with mediators for eosinophilic inflammation. Strain-level detection uncovered three non-typeable *H. influenzae* strains PittEE, PittGG and 86-028NP in the airway microbiome, where PittGG and 86-028NP abundances may inversely predict eosinophilic inflammation. The full-length 16S data augmented the power of functional inference and led to the unique identification of butyrate-producing and nitrate reduction pathways as significantly depleted in COPD.

Our analysis uncovered substantial intra-genus heterogeneity in the airway microbiome associated with inflammatory phenotypes and could be of clinical importance, thus enabled a refined view of the airway microbiome in COPD.

**“Take-home” message:** The species-level analysis using the ‘third-generation’ sequencing enabled a refined view of the airway microbiome and its relationship with clinical outcome and inflammatory phenotype in COPD.

## Introduction

The airway microbiome in chronic obstructive pulmonary disease (COPD) has been well studied in the last decade. The airway microbiome differs between health and COPD[1–3], shifts during exacerbations[4–6], associates with airway inflammation[5, 7] and predicts 1-year mortality of hospitalized exacerbation patients[8], all suggesting the implication of airway microbiome in COPD pathogenesis. Despite advances, the precise role of airway microbiome in COPD remains incompletely understood. An important knowledge gap is that our current view of airway microbiome is limited at most to its composition at the genus-level, due to insufficient resolution of one or few hypervariable regions of 16S rRNA gene being sequenced in essentially all previous studies. In these studies, certain bacterial genera were often reported to be altered as a whole in disease and in relation to airway inflammation[5, 6]. However, from an ecological perspective, members of microbial community do not necessarily function according to their taxonomic groups, instead diversified species can act in ecological “guilds” that co-adapt to altered environment[9, 10]. Therefore, the aggregated genus-level associations can be spurious or even misleading due to violation of basic ecological concepts. The inadequate depth of taxonomic profiling limits not only the accuracy of ecological inferences but also the ability to identify key bacterial species to use in follow-up experimental studies.

The recently advanced ‘third-generation’ sequencing technologies such as Pacific Biosciences (PacBio) and Nanopore is increasingly applied to microbiome studies[11, 12]. By generating long reads that extend tens of thousands of nucleotides, they offer the promise of increased taxonomic resolution by sequencing the full-length of 16S rRNA gene[13]. In these applications, the 16S amplicon is circularized and read through multiple passes before circular consensus sequences (CCS) is reported, which greatly reduced the initial high error rate (~10%) of the long-read sequencing to that comparable to short-read sequencing (~0.5%)[14, 15]. Recent development of sophisticated denoising algorithms further enable accurate bacterial species identification at single-nucleotide resolution with near-zero error rate[16]. In some situations, strain-level identity can be further resolved utilizing information on the full complement of 16S rRNA gene alleles in bacterial genomes[16, 17].

Here we report the first comprehensive analysis of airway microbiome in COPD at species-level using PacBio sequencing. We also attempted to resolve strain-level identity when possible. Our results showed that there was substantial intra-genus diversity and heterogeneity in the airway microbiome that was previously underappreciated, which was associated with patient clinical features and airway inflammatory phenotypes.

## Methods

### Subjects and samples

Sputum samples of 98 stable COPD patients and 27 age-matched healthy controls were collected in the First Affiliated Hospital of Guangzhou Medical University. The study was approved by the ethics committee of the First Affiliated Hospital of Guangzhou Medical University (No. 2017-22) and was registered in www.clinicaltrials.gov (NCT 03240315). All COPD patients met the diagnostic criteria according to GOLD guideline and were assessed for symptoms and exacerbation frequency (Table 1). Patients with antibiotic usage within 4 weeks were excluded. Induced sputum were obtained for all subjects and quality-controlled. A panel of 47 sputum mediators were measured in a subset of 59 patients using custom antibody microarray[18]. Additional information on sequencing, reagent controls, qPCR, and statistical analyses are provided in the supplementary document.

**Table 1.** Major demographic and clinical characteristics of subjects.

| Demographic and clinical features | Healthy<br>(n=27) | COPD<br>(n=98) | P-value |
| --- | --- | --- | --- |
| Age | 65.4 (10.8) | 66.2 (8.9) | 0.67 |
| Gender, n(M/F) | 23/4 | 89/9 | 0.48 |
| Current smoking, n(Y/N) | 9/18 | 85/13 | 1.0e-4*** |
| GOLD (1/2/3/4) | NA | 24/33/32/9 | NA |
| New GOLD (a/b/c/d)<sup>\$</sup> | NA | 39/38/4/17 | NA |
| Frequent exacerbator (Y/N) <sup>\$\$</sup> | NA | 20/78 | NA |
| ICS usage (Y/N) | NA | 58/40 | NA |
| pre-FEV<sub>1</sub> (L) | 2.8±0.1 | 1.5±0.1 | 5.1e-10*** |
| pre-FVC (L) | 3.4±0.2 | 2.9±0.1 | 0.01** |
| pre-FEV<sub>1</sub> (%) | 100.0±2.6 | 56.1±2.7 | 1.0e-10*** |
| pre-FEV<sub>1</sub>/FVC | 0.81±0.01 | 0.49±0.02 | 2.5e-13*** |
| post-FEV<sub>1</sub> (L) | NA | 1.6±0.7 | NA |
| post-FVC (L) | NA | 3.1±0.8 | NA |
| post-FEV<sub>1</sub> (%) | NA | 59.6±2.6 | NA |
| post-FEV<sub>1</sub>/FVC | NA | 0.5±0.1 | NA |
| CAT score | NA | 4.0±0.6 | NA |
| mMRC | NA | 0.4±0.1 | NA |
| Total sputum cells (cells×10<sup>9</sup>/L) | NA | 20.4±2.8 | NA |
| Sputum neutrophils (%) | NA | 86.2±1.2 | NA |
| Sputum eosinophils (%) | NA | 5.3±0.7 | NA |
| Sputum lymphocyte (%) | NA | 0.7±0.1 | NA |
| Sputum monocyte (%) | NA | 7.9±1.1 | NA |

Continuous data are present as mean (range) or mean±SEM.
P-value was calculated using Fisher exact test for categorical variables and using Wilcoxon rank-sum test for continuous variables. \*\*\* P<0.001; \*\* P<0.01; \* P<0.05
ICS: inhaled corticosteroids; FEV<sub>1</sub>: forced expiratory volume in one second; FVC: forced vital capacity, CAT: COPD Assessment Test, mMRC: modified Medical Research Council.
<sup>\$</sup> The new GOLD classification based on mMRC, CAT and exacerbation frequency[25].
<sup>\$\$</sup> The frequent exacerbator was defined as exacerbation event ≥2/last year.

### PacBio sequencing and analysis

Bacterial genomic DNA was extracted from selected sputum plugs using Qiagen DNA Mini kit. Negative controls for extraction and PCR were sequenced with all samples. The full-length 16S rRNA gene was amplified using barcoded 27F and 1492R primers and sequenced using PacBio Sequel. Circular consensus sequences (CCS) were generated using the *ccs* application in SMRTLink 5.1 with minPasses=5 and minPredictedAccuracy=0.90. The demultiplexed CCS were analyzed using DADA2 v1.12.1 recently customized for the PacBio full-length 16S sequencing data[16, 19]. Amplicon sequence variants (ASVs) were assigned to species only if they had unique, 100% identity match to a single species. Sequences were rarefied to 3,119 reads (Figure S1).

### Strain-level identification

Callahan et al. described a method for strain-level identification using full-length 16S data leveraging the full complement of 16S rRNA alleles in bacterial genomes[16]. In principle, a strain can be confidently assigned if all intra-genomic 16S sequence variants of that strain are recovered in integral ratios according to its genuine allelic variants. In extension to this approach, we designed a pipeline to assign strain-level bins in three steps. 1) All species-level ASVs were BLASTn-searched against NCBI-nt database. ASVs with 100% identity to the same bacterial genome were assigned to the same initial bins. 2) The ASVs within each initial bin were subject to pairwise Pearson correlation, to generate refined bins by identifying ASVs with co-occurrence pattern (Pearson’s R>0.7). 3) For each refined bins, the copy number ratio of ASVs were determined based on linear regression coefficient, and reconciled with the genuine copy number ratio of the 16S alleles in the corresponding bacterial genomes. The ASVs in integral copy number ratio with the genuine ratio were retained in the final bins and assigned with strain-level taxonomy.

### Statistical analysis

Differential microbiome features between COPD and controls were identified using linear discriminant analysis (LDA) effect size (LEfSe) method with LDA>2.0[20]. Random forest analysis was performed using Weka with 7-fold cross-validation[21]. To identify microbiome-mediator associations independent of patient demographic factors, all microbiome features and the 47 sputum mediators were first residualized using a general linear model adjusting for covariates such as age, gender and smoking history. An all-against-all correlation analysis was performed on the residues of microbiome features and mediators using HAllA[22], and was subject to unsupervised clustering. Co-occurrence analysis of microbiome was performed using SparCC[23]. Functional inference of microbiome was performed using PICRUSt2[24]. The false discovery rate (FDR) method was used to adjust *P*-values.

## Results

### Overview of the species-level airway microbiome profile

A total of 2,635,140 high-quality CCS reads were obtained for 98 stable COPD patients and 27 controls (Table 1). The average number of passes on the 16S gene was 34.9 for all CCS, equivalent to a low error rate of ~0.48% based on previous sequencing runs on a mock community[17]. A total of 2,868 non-singleton ASVs were identified, of which 795 ASVs (27.7%) were putatively assigned to 228 bacterial species from 92 genera. Twenty species had an average relative abundance greater than 0.005 (Table 2). The number of species capable of being detected increased by 3.26 folds compared to a re-analysis of all previous COPD airway microbiome studies using the same pipeline (Table S1). *Streptococcus*, *Prevotella* and *Neisseria* had the highest number species identified (Figure S2a). There was significant community shift in COPD versus controls (Figure 1a-b, Adonis, *P*=0.004). LEfSe analysis identified 11 discriminatory species between COPD and controls (Figure 1c, LDA>2.0). Random forest analysis using these 11 species yielded significantly increased precision in classifying patients, compared to that using 9 discriminatory genera with the same criteria (LDA>2.0) (Figure 1d, Figure S2b, AUC: 0.787 versus 0.706, *P*=0.026). Figure 1e showed an overview of species-level airway microbiome profile.

**Table 2.** The major species-level taxa identified in this study (average relative abundance>0.005).

| Species | All<br>average | COPD<br>average | Healthy<br>average | Fold<br>change | FDR<br>P-value |
| --- | --- | --- | --- | --- | --- |
| *Prevotella intermedia* | 0.005 | 0.006 | 0.005 | 1.191 | 0.319 |
| *Prevotella melaninogenica* | 0.060 | 0.057 | 0.071 | 0.801 | 0.342 |
| *Prevotella pallens* | 0.013 | 0.012 | 0.017 | 0.700 | 0.0140* |
| *Streptococcus*<br>*pseudopneumoniae* | 0.017 | 0.020 | 0.004 | 5.667 | 0.222 |
| *Streptococcus salivarius* | 0.013 | 0.015 | 0.004 | 3.664 | 0.277 |
| *Streptococcus thermophilus* | 0.027 | 0.030 | 0.014 | 2.228 | 0.809 |
| *Haemophilus influenzae* | 0.043 | 0.052 | 0.008 | 6.779 | 0.617 |
| *Haemophilus*<br>*parahaemolyticus* | 0.009 | 0.010 | 0.004 | 2.564 | 0.204 |
| *Haemophilus parainfluenzae* | 0.012 | 0.009 | 0.020 | 0.474 | 0.0074** |
| *Neisseria meningitidis* | 0.009 | 0.009 | 0.008 | 1.044 | 0.128 |
| *Neisseria mucosa* | 0.007 | 0.008 | 0.002 | 4.452 | 0.785 |
| *Neisseria perflava* | 0.011 | 0.012 | 0.005 | 2.391 | 0.742 |
| *Neisseria subflava* | 0.017 | 0.012 | 0.036 | 0.330 | 0.0415* |
| *Fusobacterium nucleatum* | 0.016 | 0.013 | 0.027 | 0.495 | 0.139 |
| *Fusobacterium*<br>*periodonticum* | 0.014 | 0.012 | 0.022 | 0.531 | 0.0199* |
| *Pseudomonas aeruginosa* | 0.016 | 0.020 | 0.000 | NA | 0.105 |
| *Moraxella catarrhalis* | 0.030 | 0.038 | 0.001 | 28.947 | 0.275 |
| *Veillonella parvula* | 0.005 | 0.004 | 0.010 | 0.439 | 0.208 |
| *Porphyromonas gingivalis* | 0.010 | 0.009 | 0.015 | 0.570 | 0.282 |
| *Campylobacter concisus* | 0.006 | 0.006 | 0.007 | 0.902 | 0.233 |

P-value was calculated using Wilcoxon rank-sum test. \*\*\* FDR $P < 0.001$ ; \*\* $P < 0.01$ ; \* $P < 0.05$

**Figure 1.**
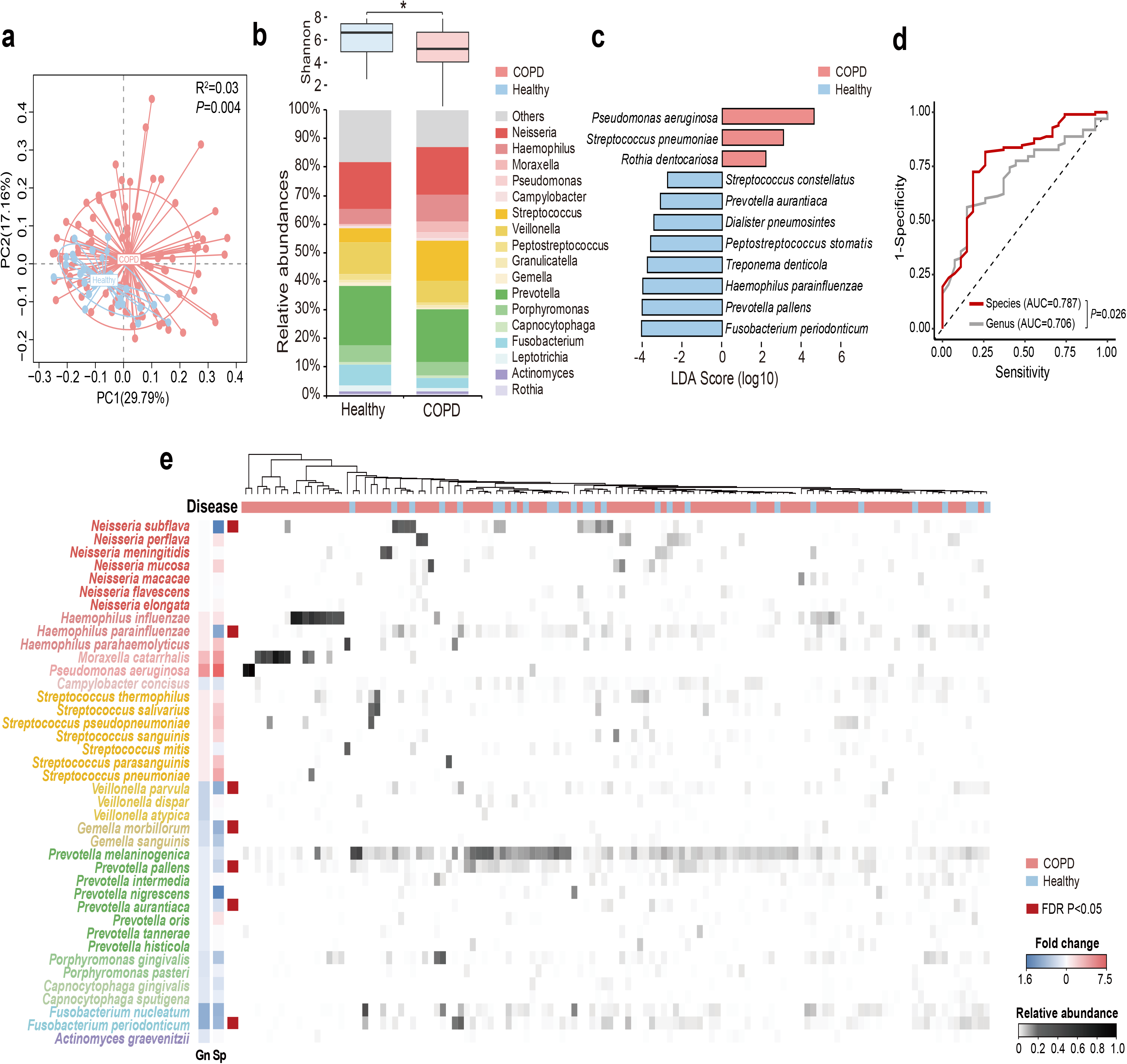
The overview of species-level profile of the airway microbiome in COPD patients and healthy controls. **a)** Principal coordinate analysis based on weighted UniFrac distance on sputum samples from 98 COPD patients and 27 healthy controls. **b)** The Shannon diversity and relative abundances of major genera (relative abundance>0.005) in COPD patients and healthy controls. **c)** The 11 top discriminatory species-level taxa between COPD and controls as identified from LEfSe analysis (LDA>2.0). **d)** The receiver operating characteristic curves for the Random Forest analyses using the 11 species-level and 9 genus-level discriminatory taxa (LDA>2.0) to segregate COPD patients from controls. **e)** The heatmap for the species-level microbiome profile. The major species-level taxa (relative abundance>0.001) within each genus in panel b) were shown. The fold change of each species (Sp) and its corresponding genus (Gn) in COPD patients versus controls were shown beside the taxonomy.

Overall there were no significant microbial community shifts between smokers and non-smokers within COPD patients or healthy controls, between patients with and without inhaled corticosteroid usage, and between frequent and non-frequent exacerbators (defined as exacerbation events>=2/last year, Figure S3a-b). Among species with relative abundance>0.001, *Haemophilus parahaemolyticus* was significantly increased in COPD smokers (Fold-change=6.40, FDR *P*=0.02, Figure S3c). *Ralstonia mannitolilytica*, an opportunistic pathogen, was significantly increased in frequent exacerbators (Fold-change=4.94, FDR *P*=0.005, Figure S3c). The increase of *R. mannitolilytica* was further confirmed by qPCR (Figure S3d).

### Substantial intra-genus heterogeneity in the airway microbiome

Inspection of individual species revealed substantial intra-genus heterogeneity in their relationships with COPD. For example, while *Neisseria mucosa* was increased in COPD versus controls, its counterpart *Neisseria subflava* was significantly depleted (Figure 1a). The reciprocal relationships with COPD were also observed between *Haemophilus influenzae* and *Haemophilus parainfluenzae*, and between *Prevotella oris* and other *Prevotella* species (Figure S4). The species also altered differently with enhanced disease severity. For example, *H. parainfluenzae* and *N. subflava* were the most predominant species within the respective genera in healthy subjects, while *H. influenzae* and *N. meningtidis* took over and became over-dominant in GOLD IV patients (Figure 2a). Within *Streptococcus*, *Streptococcus salivarius* and *Streptococcus thermophilus* were most highly abundant in GOLD I patients, whereas *Streptococcus pseudopneumoniae* and *Streptococcus pneumoniae* became dominant in GOLD II and IV patients respectively (Figure 2a). Similar turnovers were also observed in patients classified using new GOLD classification scheme based on mMRC, CAT score and exacerbation frequency[25] (Figure S5). Opposite relationships with patient sputum neutrophilic levels were further observed between *H. influenzae* and *H. parainfluenzae* (Figure 2b), and between *Prevotella melaninogenica* and *Prevotella denticola* (Figure S6a). Individual species within the same genus exhibited disproportionately more co-exclusive than co-occurrence relationships (Figure 2c, Figure S6b), indicating ecological competition. qPCR using primers designed on species-specific genes showed concordance between the absolute count and relative abundance of *H. influenzae* and *H. parainfluenzae* (Figure 2d), as well as two other paired species within *Streptococcus* and *Prevotella* (Figure S6c), indicating accuracy of our approach in species quantification. These results suggested that there were substantial intra-genus heterogeneity resulting from interspecific competition in the airways.

**Figure 2.**
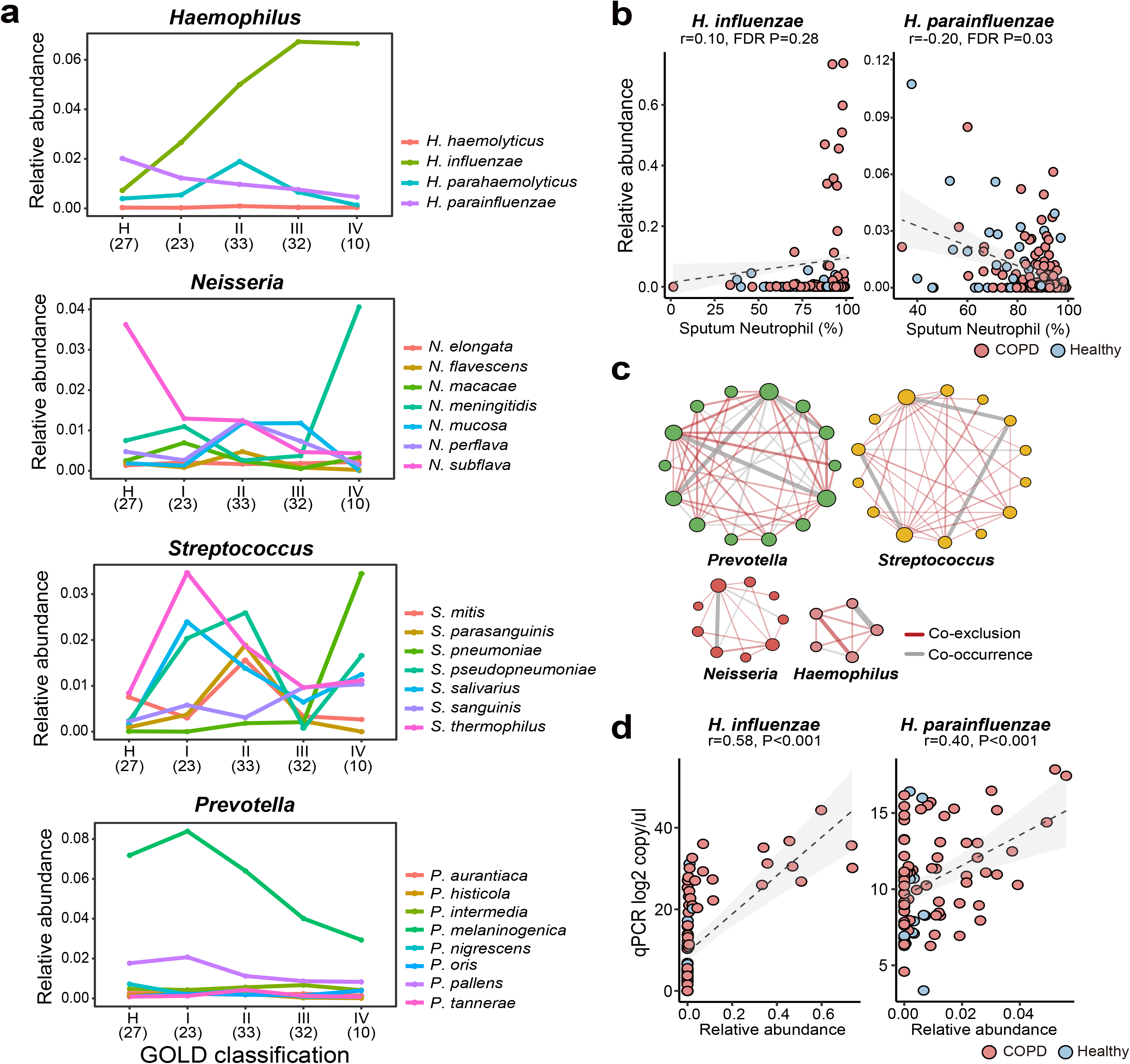
The intra-genus heterogeneity of the airway microbiome. **a)** The alternation of major species in *Haemophilus*, *Neisseria*, *Streptococcus* and *Prevotella* between healthy controls and COPD patients with increasing disease severity based on GOLD classification (spirometry-based). The number of subjects in each subgroup was indicated in the parenthesis. **b)** The reciprocal relationship between *H. influenzae* and *H. parainfluenzae* with sputum neutrophilic percentage. **c)** More pervasive co-exclusive than co-occurrence relationships between major species in *Prevotella*, *Streptococcus*, *Neisseria* and *Haemophilus*. Only significant correlations were shown in the networks (SparCC, *P*<0.05). Co-exclusion relationships were in red, whereas co-occurrence relationships were in grey. **d)** qPCR assays using species-specific primers showed concordance between absolute counts and relative abundances of *H. influenzae* and *H. parainfluenzae*.

### Specific bacterial species were associated with neutrophilic or eosinophilic inflammation

To investigate how the intra-genus heterogeneity was related to airway inflammation, we performed an all-against-all correlation analysis between the species-level microbiome features and a panel of 47 sputum inflammatory mediators measured in a subset of 59 COPD patients. We used residualized correlation to identify microbiome-mediator correlations independent of patient demographic co-factors[22]. Unsupervised clustering based on the correlation profile revealed four clusters of bacterial species that each had distinct association patterns with three groups of mediators (Group 1-3, Figure 3). Four pathogens, *Moraxella catarrhalis, Pseudomonas aeruginosa, N. meningtidis* and *H. influenza*e, exhibited negative associations with a group of 11 mediators mostly Th2-related (i.e. IL-5, IL-13, CCL17), while they were positively correlated with a group of 21 mediators mostly Th1, Th17-related or pro-inflammatory (i.e. IL-8, IL-17, MMP-8), and had mixed relationships with the remaining mediators. By contrast, another seven species, *Prevotella aurantiaca, Fusobacterium nucleatum, Leptotrichia buccalis, Prevotella histicola, Porphyromonas gingivalis, N. mucosa* and *Tropheryma whipplei*, were specifically associated with increased Th2 mediators. Members of the two groups of mediators further showed specific correlations with increased sputum neutrophil or eosinophil percentages respectively (FDR *P*<0.05), in agreement with their roles in neutrophilic or eosinophilic inflammation. Correspondingly, all seven species were increased in the eosinophilic COPD patients (eosinophil>3%, Figure S7a). The increase of *T. whipplei* was further confirmed by qPCR (Figure S7b). Such clustering patterns were however not observed at the genus-level (Figure S8), indicating microbiome associates with airway inflammatory phenotypes in a species-specific manner.

**Figure 3.**
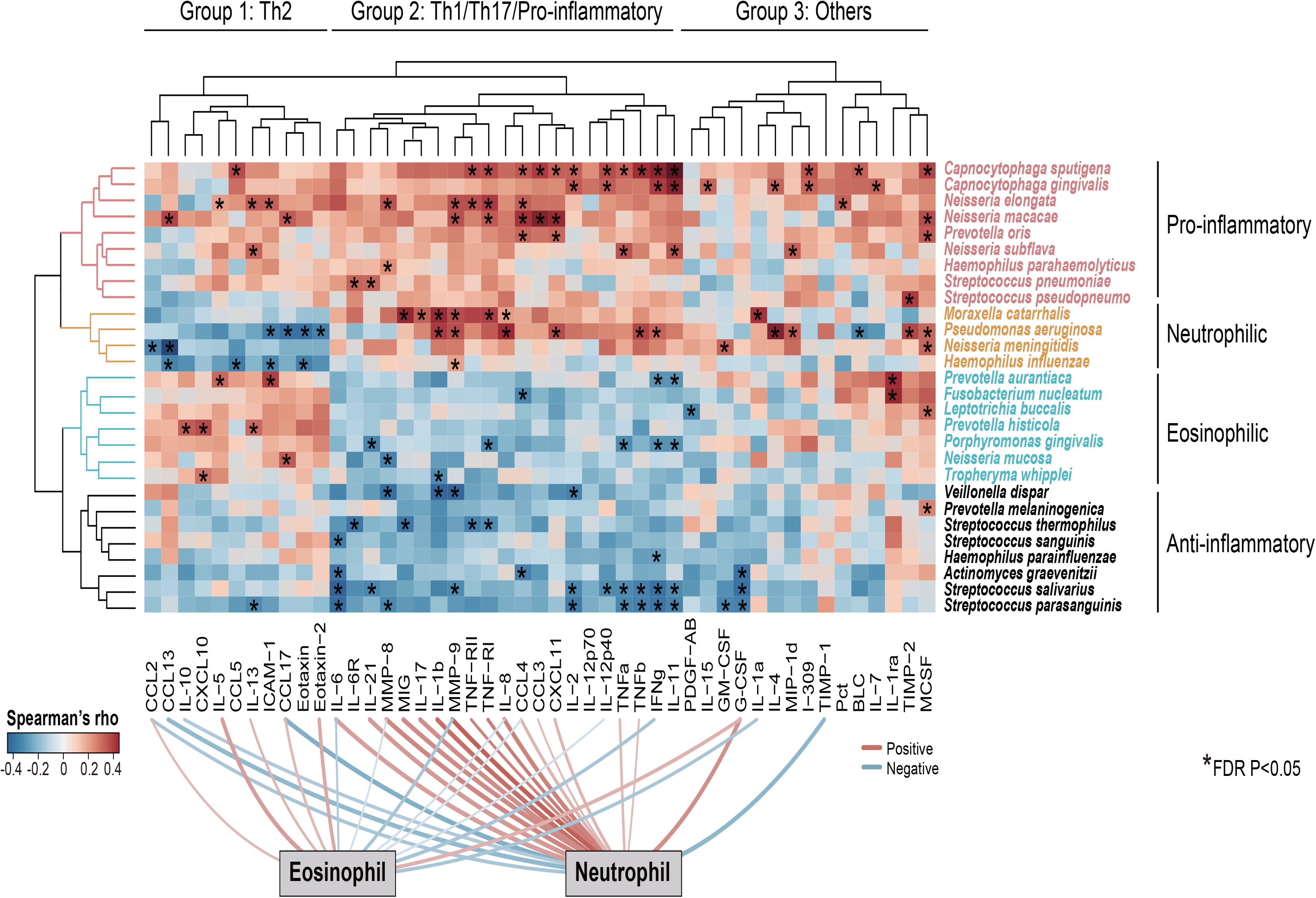
Species-specific association of airway microbiome with inflammatory phenotypes. Unsupervised hierarchical cluster analysis on an all-against-all correlation profile between species-level microbiome features and 47 sputum mediators from a subset of 59 COPD patients (Ward’s method). The species were shown if they had relative abundance>0.001 and were significantly associated with at least one of the 47 sputum mediators (HAllA, FDR *P*<0.05). The mediators were clustered into three groups and termed based on their classes and associations with airway eosinophils or neutrophils (Group 1: Th2-related, Group 2:Th1/Th17/Pro-inflammatory-related, Group 3: Others). The microbiome features were clustered into four groups based on their association patterns with the three groups of mediators (termed “Pro-inflammatory”, “Neutrophilic”, “Eosinophilic” and “Anti-inflammatory”). The significant associations were indicated in asterisks. Significant positive and negative associations between sputum mediators and neutrophilic and eosinophilic percentages were shown on bottom of the heatmap (FDR *P*<0.05).

### Strain-level identification of the airway microbiome

We further explored possible strain-level diversity in the airway microbiome. Recent studies showed that it is possible to resolve strain-level identity using full-length 16S sequences by leveraging the power of the full complement of 16S rRNA alleles within bacterial genomes[16, 17]. Using a set of stringent criteria (see methods, Figure S9), we were able to identify ASV bins corresponding to 10 bacterial strains (Table S2). For the first time, we identified three non-typeable *H. influenzae* (NTHi) strains PittEE, PittGG and 86-028NP in the airway microbiome, although the major allele of 86-028NP was not detected (Figure 4a-b). All three strains increased in COPD versus controls, and were associated with distinct groups of mediators (Figure 4c). Notably, 86-028NP and PittGG exhibited inverse associations with Th2 chemokines such as CCL17 and CCL13 related to eosinophilic inflammation. qPCR using strain-specific primers validated our results in PittEE and PittGG (Figure 4d), although the strain detection rate by sequencing was lower than that using qPCR. qPCR for 86-028NP yield positive but non-significant correlation (Figure 4d).

**Figure 4.**
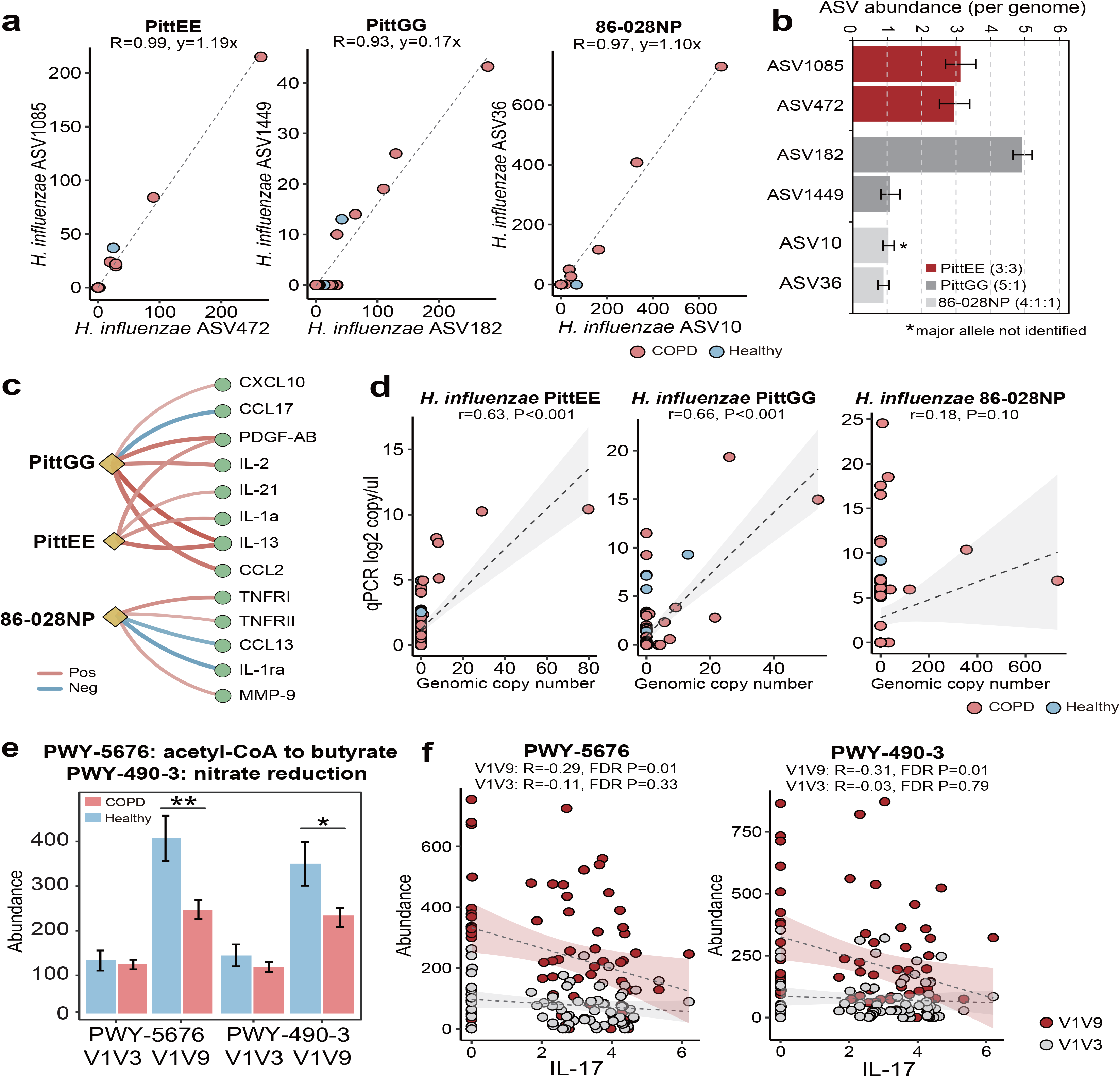
Strain-level identification and functional inference of the airway microbiome. **a)** The strong correlation pattern between pairs of ASVs assigned to the strains PittEE, PittGG and 86-028NP (Pearson’s *R*>0.93). **b)** The copy number of the highly-correlated ASVs are in integral ratio with the genuine allelic frequency of the 16S rRNA genes within the genome (PittEE 3:3, PittGG 5:1, 86-028NP 4:1:1), supporting the assignment of the ASVs to the corresponding strains. The major 16S allele of the 86-028NP strain was not detected. **c)** Significant associations between the three *H. influenzae* strains with sputum mediators (Spearman, FDR *P*<0.05). **d)** qPCR results using strain-specific primers for the three *H. influenzae* strains in relation to their relative abundances in the sequencing data. **e)** The abundances of the two pathways ‘PWY-5676: acetyl-CoA fermentation to butyrate’ and ‘PWY-490-3: nitrate reduction’ in COPD and healthy subjects as inferred from the full-length (V1V9) and V1V3 data using PICRUSt2 (** FDR *P*<0.01, * FDR *P*<0.05). **f)** The two pathways showed negative correlations with IL-17, which was more pronounced when inferred from full-length 16S sequences than V1V3 sequences.

### A systematic evaluation of 16S sub-regions for airway microbiome profiling

The full-length 16S sequences can serve as a benchmark for a systematic evaluation on the performance of individual hypervariable regions for airway microbiome studies. We created partitions of 16S sequences from the full-length data according to nine hypervariable regions used in previous COPD microbiome studies, and analyzed each partition separately. Among all sub-regions, V1V3 and V3V4 were the highest in the number of species assigned as well as the proportion of sequences assigned to species (Figure S10a). In addition, the V1V3 and V3V4 regions captured the greatest microbial beta diversity measured using pairwise Bray-Curtis dissimilarity, whereas the diversity was the lowest for V4 (Figure S10a). The V4 region was particularly poor in classifying Proteobacteria and Actinobacteria, with 79.8% and 90.9% of sequences from these two phyla unable to be assigned to species (Figure S10b). The V1V3 region also bear the highest similarity with the full-length data in microbial community composition (Mantel test, Figure S10c).

### Full-length 16S sequences enhanced the power of functional inference

PICRUSt is a useful tool to infer functional capacity of microbiome based on 16S sequences[26]. PICRUSt analysis using the full-length 16S data enhanced the power of functional inference by increasing the predicted pathway abundances by an average 1.83 fold compared to individual sub-regions. Again, V1V3 were next best in terms of the predicted pathway abundances (Figure S10a).

The augmented power of functional prediction led to the unique identification of 9 pathways as disease-associated using the full-length 16S data (Table S3). Of interest are two pathways ‘acetyl-CoA fermentation to butyrate’ and ‘nitrate reduction’, both inferred as significantly depleted in COPD (FDR *P*<0.05, Figure 4e, Figure S11). qPCR using validated broad-spectrum primers on butyryl-CoA:acetate-CoA-transferase gene[27] in the butyrate pathway confirmed our findings by showing 4.32 fold decrease of the gene in COPD versus controls (Table S4). Furthermore, the two pathways showed inverse correlations with IL-17, which were more pronounced when inferred from full-length 16S data than from sub-regions (Figure 4f).

## Discussion

Here we provided the first comprehensive insights on the COPD airway microbiome at the species and strain-levels. By applying the ‘third-generation’ sequencing to the full-length 16S rRNA gene, we uncovered diversity and complexity in the airway microbiome at in-depth taxonomic levels that were previously underappreciated. In light of our results, many aspects of our understanding of the COPD airway microbiome need to be refined.

Our results showed that there were substantial intra-genus heterogeneity in the airway microbiome in relation to patient clinical outcomes. Individual species within the same genus often altered differentially in COPD and with enhanced clinical severity and exacerbation frequency. The species predominant in healthy state can be taken over by another species within the same genus in severe COPD patients. Therefore, the genus-level associations reported in all previous airway microbiome studies likely represent a weakened signal confounded by the mixed effects of individual species within and should therefore be interpreted with caution. Unsupervised cluster analysis identified two groups of bacterial species showing specific associations with mediators related to neutrophilic or eosinophilic inflammation respectively. The neutrophil-associated species included respiratory pathogens like *H. influenza*e and *Moraxella catarrhalis*[28]. The eosinophil-associated species included *T. whipplei*, a clinically important species reported to be implicated in pneumonia[29], HIV infection[30] and eosinophilic, corticosteroid-resistant asthma[31, 32]. Such clustering pattern was non-existent at the genus-level. Hence the species-level delineation enabled a more ecologically coherent view of airway microbiome according to inflammatory phenotypes.

In extension to a previous approach[16], we detected three NTHi clinical strains PittEE, PittGG and 86-028NP in the airway microbiome with reasonably high confidence, based on which the strain-level heterogeneity was also observed in the airways. All three strains were initially isolated from otitis media patients[33–35]. It has been shown that the PittGG strain, by possessing an extra cluster of 339 genes and a Hif-type pili structure, conveyed greater virulence than PittEE[34]. qPCR assays based on *alpA* gene on this extra locus confirmed our results in PittGG quantification. While all three strains were related to increased Th1/Th17 mediators, 86-028NP and PittGG were further associated with decreased Th2-related CCL13 and CCL17, indicating their abundances may negatively predict eosinophilic inflammation. We realize that the strain-level diversity and detection rate remained relatively low, which is a caveat due to inherently limited power of 16S sequences in strain-level resolution and its sensitivity to potential sequencing errors.

We identified *Ralstonia mannitolilytica* as significantly increased in COPD patients with frequent exacerbator phenotype. *R. mannitolilytica* is an opportunistic pathogen that has been recovered from cystic fibrosis airways[36]. In a previous report, the same species was isolated from one COPD exacerbation patient in western China with extreme symptoms and acute respiratory failure[37]. *Ralstonia* spp. rarely cause infection in healthy individuals but can be a severe pathogen especially in immunosuppressed patients[38]. Therefore, the presence of *R. mannitolilytica* in stable COPD patients may be an important contributing factor in predisposing patients to recurrent infection and exacerbations.

The systematic comparison of 16S sub-regions indicated that V1V3 performed the best in terms of microbial diversity and the power of functional inference. Our results are consistent with the analysis by Johnston et al.[17], and should guide future studies that sequencing V1V3 region may be a surrogate for the full-length 16S data. Conversely, sequencing V4 alone, despite its wide usage in airway microbiome studies, might not provide sufficient resolution for in-depth taxonomic profiling.

With augmented power in functional inference, we identified butyrate-producing and nitrate reduction pathways as uniquely depleted in COPD using full-length 16S data. Butyrate is a well-characterized microbial metabolite with anti-inflammatory effects[39], and nitric oxide, the end product of nitrate reduction, may also have disease-ameliorating role via suppressing NLRP3 inflammasome activation[40]. Functional validations are warranted to explore these microbial metabolites as novel therapies for COPD.

The limitations of this study include its single-centered, cross-sectional design, the relatively small group of healthy subjects, and the absence of sufficient data to explore species-specific relationships with other etiological factors such as viral infections. The species-level characterization on larger, longitudinal cohorts is necessary to understand how species alter differentially during exacerbations and to treatment, the temporal dynamics of ecological heterogeneity, and its underlying relationships with airway inflammation and disease progression.

In summary, we reported the comprehensive landscape of COPD airway microbiome at species and strain-levels. We showed there was substantial intra-genus heterogeneity associated with patient clinical outcome and inflammatory phenotypes. Sequencing the full-length 16S rRNA gene enabled a refined, ecologically coherent view on the composition and function of the COPD airway microbiome, and should see a wider applicability in airway microbiome studies in future.

## Acknowledgement

This work was supported by the National Key R%D Program of China (2017YFC1310600) funded to HZ and RC, and the National Natural Science Foundation of China (31970112) funded to ZW.

