## Supplementary document for "A Refined View of Airway Microbiome in Chronic Obstructive Pulmonary Disease at Species and Strain-levels"

**Supplementary Methods**

**Patients inclusion and exclusion criteria**

For COPD patients, the inclusion criteria were: 1) age >40 years; and 2) confirmed diagnosis of COPD according to the GOLD guideline[1] (post-bronchodilator forced expiratory volume in 1s [FEV1]/forced vital capacity [FVC] ratio <0.7). The exclusion criteria were: 1) diagnosis of known respiratory disorders other than COPD; 2) COPD exacerbation within 4 weeks of enrolment; 3) history of lung surgery and tuberculosis; 4) diagnosis of cancer; 5) blood transfusion within 4 weeks of enrolment; 6) diagnosis of autoimmune diseases; 7) enrolment in a blinded drug trial; and 8) antibiotic usage within 4 weeks of enrolment. Informed consent was obtained from all patients.

**Quality control of sputum samples**

All sputum samples were subjected to quality control upon collection. Briefly, sputum plugs which contained the most viscous material were picked up in a petri dish and isolated from saliva. The selected sputum plugs were prepared for cytology by dilution with 0.1% dithiothreitol (DTT) solution and filtered through 48 µm nylon-mesh filter according to standardized sputum processing protocol[2]. The numbers of total cells, squamous epithelial cells and leukocytes were counted and recorded. Sputum specimens with squamous epithelial cells:leukocytes < 1:2.5 were considered unlikely to be contaminated with oropharyngeal flora and acceptable for downstream experiments[3, 4].

**Sputum cytokines and chemokines**

Sputum supernatant were collected from a subset of 59 COPD patients with sufficient sputum samples, using the procedures described above. A panel of 47 sputum cytokines and chemokines (BLC, Eotaxin, Eotaxin-2, CXCL11, CXCL10, CCL2, CCL3, CCL4, CCL5, CCL13, CCL17, G-CSF, GM-CSF, I-309, ICAM-1, IFNg, IL-1a, IL-1ra, IL-1b, IL-2, IL-4, IL-5, IL-6, IL-6R, IL-7, IL-8, IL-10, IL-12p40, IL-12p70, IL-13, IL-15, IL-16, IL-17, IL-21, MCSF, MIG, MIP-1d, MMP-8, MMP-9, PDGF-AB, Procalcitonin, TNFa, TNFb, TNFRI, TNFRII, TIMP-1, TIMP-2) were measured from sputum supernatant in quadruplicate, using a customized microarray[5] (Human Cytokine Antibody Microarray slides; RayBiotech Inc., Norcross, GA, USA).

**DNA extraction and sequencing**

Bacterial genomic DNA was extracted from selected sputum plugs using Qiagen DNA Mini kit as per the manufacturer’s instruction. Negative controls for extraction (no sputum) and PCR amplification (no DNA template, ddH2O only) were included in each experiment. The extraction negative controls were subsequently sequenced together with all samples to identify any potential contaminating bacterial species. The full-length (V1V9) bacterial 16S rRNA gene sequences were amplified using barcoded 27F (AGRGTTYGATYMTGGCTCAG) and 1492R (RGYTACCTTGTTACGACTT) primers. Library construction was performed using Pacific Biosciences (PacBio) SMRTbell™ Template Prep Kit V1 on normalized pooled PCR products. Sequencing was performed using Pacific Biosciences (PacBio) Sequel platform.

**Sequencing reagent controls**

Reagent controls for extraction (no sputum material) and PCR amplification (no template, ddH_2_O only) were included in the experiment. One DNA extraction controls and one PCR negative control were subsequently sequenced to identify any potential contaminating bacterial species.

The Amplicon Sequence Variants (ASVs), their number of sequences and taxonomy in the reagent controls were presented in **Table S5**. None of the bacterial ASVs present in the reagent controls have >30 read counts and none of them were the major members in the airway microbiome. Although negative reagent controls were performed for DNA extraction and PCR amplification step, we performed a retrospective analysis to ensure that potential contamination risks were minimized. We compared our results against the 92 contaminant genera detected in sequenced negative ‘blank’ controls by Salter et al[6]. We failed to detect 62 out of the 92 contaminant genera in our dataset (**Table S6**). Of the remaining genera that were found in our data, none had an average relative abundance greater than 8.43e-4, or had a relative abundance greater than 0.01 in any particular sample, except for *Streptococcus* which is a known member in sputum microbiome, and *Ralstonia*, *Stenotrophomonas*, *Acinetobacter* and *Pseudomonas* which were respiratory pathogens present only in a small number of patients and therefore not considered to be reagent contaminations.

**Sequence processing and analysis**

Raw reads were demultiplexed using the *lima* application, specifying that the same barcodes were attached at both ends of an insert using the flags *same* and *peek-guess*. Circular consensus sequence (CCS) reads were generated using the *ccs* application in the SMRTLink 5.1 software with parameters minPasses=5 and minPredictedAccuracy=0.90, according to an established protocol for clinical samples[7].

The demultiplexed CCS reads were quality-controlled and analyzed using the most recent DADA2 program customized for the PacBio full-length 16S rRNA sequencing data (<https://benjjneb.github.io/LRASManuscript/LRASms_fecal.html>). Briefly, primers were first removed from the CCS sequences using the *removePrimers* function. Sequences without primer matches were discarded. The remaining CCS sequences were then filtered using the *filterAndTrim* function with parameters: minLen=1000, maxLen=1600, maxN=0, maxEE=2, and minQ=3. Sequences were then dereplicated and used for learning the dataset-specific error model using the *learnErrors* function with parameter errorEstimationFunction=dada2:::PacBioErrfun. Sequences were denoised using the error model and Amplicon Sequence Variants (ASVs) were identified using the *dada* function. Taxonomy up to the genus-level was assigned using the *assignTaxonomy* function based on the silva_nr_v128_train_set sequence database by default. Species-level taxonomy was assigned using the *addSpecies* function. ASVs were assigned to species only if they had exact match (100% identity) and unique match to a single species (all exact matches were to the same species) in the silva_species_assignment_v132 sequence database, according to the dada2 protocol (<https://benjjneb.github.io/dada2/assign.html>). Sequences were rarefied to 3,119 reads per sample, according to the lowest sequencing depth for all samples. Alpha and beta diversity analysis were then performed based on the rarified ASV table using QIIME 2.0 (version 2019.10). The raw Fastq data of this study has been deposited in the Chinese National Gene Bank (CNGB) Nucleotide Sequence Archive (CNSA) under accession code CNP0000837.

**Statistical analyses**

Differential microbiome features between COPD and healthy controls were identified using a linear discriminant analysis (LDA) effect size (LEfSe) method with a threshold of logarithmic LDA score 2.0[8]. Random forest analysis was performed using genus and species-level microbiome features selected by LEfSe (LDA>2.0) using Weka 3.8 with 7-fold cross-validation[9]. Area under receiver operative characteristic curve (AUC) were assessed to evaluate the performance of the random forest models. Statistical comparison of AUCs was performed using pROC package in R[10]. To identify microbiome-mediator associations independent of patient demographic factors, we performed a residualized correlation analysis[11]. All microbiome features and the 47 sputum mediators were first residualized using a general linear model adjusting for patient demographic covariates including age, gender, smoking history, exacerbation frequency and ICS usage. An all-against all correlation analysis was performed on the residues of microbiome features and sputum mediators using HAllA (Hierarchical All-against-All association testing)[11], a computational tool to identify pairs of statistically significant assocations in a hierarchical manner without getting tripped by the numerous multiple hypothesis testing resulting from the high dimensionality of the omics data (FDR *P*<0.05). Unsupervised hierarchical cluster analysis was performed on the correlation profile using Ward’s method. The mediators were clustered into three groups and termed based on their classes and associations with airway eosinophils or neutrophils (Group 1: Th2-related, Group 2: Th1/Th17/Pro-inflammatory-related, Group 3: Others). The microbiome features were clustered into four groups based on their association patterns with the three groups of mediators (termed “Pro-inflammatory”, “Neutrophilic”, “Eosinophilic” and “Anti-inflammatory”). The species in “Pro-inflammatory” or “Anti-inflammatory” groups had overall positive or negative associations with the broad range of Group 1-3 mediators. The species in “Neutrophilic” or “Eosinophilic” groups had specific positive associations with Group 2 or Group 1 cytokines, respectively. Co-occurrence and co-exclusion relationships of species-level microbiome features were determined using SparCC[12] (*P*<0.05). Functional inference of microbiome was performed using PICRUSt2[13].

Individual partitions of 16S rRNA gene sequences were generated by trimming the nine different hypervariable regions (V1V2, V1V3, V2, V3, V3V4, V3V5, V4, V6V8 and V6V9) of the full-length sequencing data according to established primer sets using Cutadapt v2.6[14]. The same DADA2 analysis, as described above for the full-length 16S data, was performed on each individual partitions of the 16S sequences. Mantel test was performed to assess the similarity in microbiome compositional and functional profiles generated using the full-length 16S sequences as well as using individual hypervariable regions.

**Strain-level taxonomy identification**

Recent studies showed that PacBio sequencing of the full-length 16S rRNA gene sequences can accurately resolve single-nucleotide substitutions that reflect intragenomic variations between 16S gene copies[7, 15, 16]. Such information, when properly account for, can aid in resolving strains of the same species. Callahan et al. employed a simple approach to manually create strain-level bins based on the expected integral ratios of the known copy number of all 16S rRNA genes within genomes[16]. In extension to their approach, we designed a semi-automated approach to create strain level bins (**Figure S9a**) with three steps below.

**1)** All species-level ASVs were subject to BLASTn search against a local NCBI nt database. ASVs with 100% percent identity with the same bacterial genome were assigned to the same initial bins (nitial bins). If the same set of ASVs had exact match with more than one bacterial genomes, they were assigned to each genome as one separate bin.

**2)** The ASVs within each initial bin were subject to pairwise Pearson correlation analysis, to refine the bins by identifying pairs of ASVs in the bins with strong correlation pattern across all samples (Pearson’s R>0.7, refined bins). ASV bins with no strong correlated pairs were discarded.

**3)** For each ASV bins retained, the copy number ratio of highly correlated ASVs were determined based on the linear regression coefficient, and reconciled with the genuine copy number of the 16S allele variants of the bacterial genomes to which the ASV bins were assigned (the manual step). The ASVs with copy number in approximate integral ratio (±0.2) with the genuine copy number ratio in the corresponding genomes were retained as final strain-level ASV bins and assigned with the corresponding strain-level taxonomies. The non-unique BLASTn matches (i.e. multiple genome-bins of the same set of ASVs) were resolved in this step using the genuine copy number ratio, when possible. The genomic abundances of each strain in each sample was calculated as the average abundances of ASVs normalized by their copy numbers in the genome.

**Quantitative PCR assays**

To validate our results for species identification and quantification, quantitative PCR assays were performed on a subset of 87 subjects with sufficient sputum genomic DNA (76 COPD patients and 11 controls). We designed species-specific primers for *Haemophilus influenzae, Haemophilus parainfluenzae, Streptococcus pneumoniae, Streptococcus thermophilus, Prevotella melaninogenica, Prevotella intermedia* and *Ralstonia mannitolilytica.* Species-specific genes were first identified using a BLASTp search of all protein-coding genes from each species against an in-house database of protein-coding genes from 2,764 complete bacterial genomes, to identify genes present in at least 90% of genomes of that species but absent in all other bacterial genomes. Primers were designed on the species-specific genes using Primer 3 (<http://primer3.ut.ee>). Primers with no primer-dimer detection and less than 50% identity with other locations in the genomes (BLASTn) were retained and one primer pair was selected for each species for qPCR assays. We also identified strain-specific protein coding genes for *Haemophilus influenzae* strains PittEE, PittGG and 86-028NP using similar strategies as above, and designed primers based on these genes. The primer sequences for *Trophyrema whipplei* were adopted from the study of Lozupone et al.[17], based on the *hsp65* gene sequence. To make standard curves for absolute quantification, an initial qPCR was performed on one of the sputum samples with sufficient quantity to amplify the PCR products using all primers. The PCR products were verified by Sanger sequencing and subject to standard *Escherichia coli* transformation procedure to obtain DNA templates with absolute copy numbers. The primers and their targeting genes for each species and sub-species were listed in **Table S7**.

For validation of functional inference, we performed qPCR assays on butyryl-CoA:acetate CoA-transferase gene (EC:2.8.3.8) using the validated broad-spectrum primers reported by Vital et al.[18] (but_3F: GHATYGGIGSTATGCC, but_3R: AAGTCWAAYTGWCCRCC). The universal marker *rpoB* gene was used as the internal control in the qPCR assays (F: GGYTWYGAAGTNCGHGACGTDCA, R: TGACGYTGCATGTTBGMRCCCATMA). The fold change of the *but* gene between COPD and healthy controls was calculated using the 2^-ΔΔCt approach.

All qPCR assays were performed using 96-well MicroAmp Fast Optical 96-Well Reaction Plate on the Applied Biosystems StepOnePlus^TM^ Real-Time System. The 20 µl reaction mixture contained 10 µl of SYBR® Select Master Mix (2x), 6 µl of microbial-free water, 2 µl DNA templates, and 1 µl forward and reverse primer each. The following cycling parameters were used: initial cycle of 95°C for 10 min; 40 cycles of 95°C for 15 s; 60°C for 1 min. All qPCR templates were run in duplicate. For standard curve calculation, each plate run included a decimal serial dilution of the corresponding double-stranded DNA templates as obtained above from 1E8 to 1E3 copies per µl. For qPCR of the functional gene (EC:2.8.3.8) using broad-spectrum primers, a high MgCl_2_ concentrations of 3 mM was used and thermocycling was performed as follows: 95°C for 2 min;95°C for 45 s, 54°C for 45 s, 72°C for 45 s (×40);10 min at 72°C, according to Vital et al.[18].

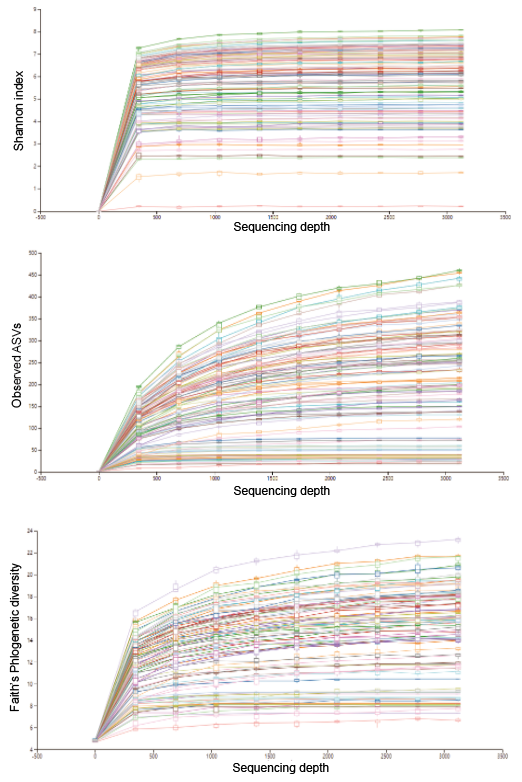

**Figure S1.** Alpha-rarefaction curve based on Shannon index, observed ASVs and Faith’s phylogenetic diversity. All samples were rarefied at the depth of 3,119 reads.

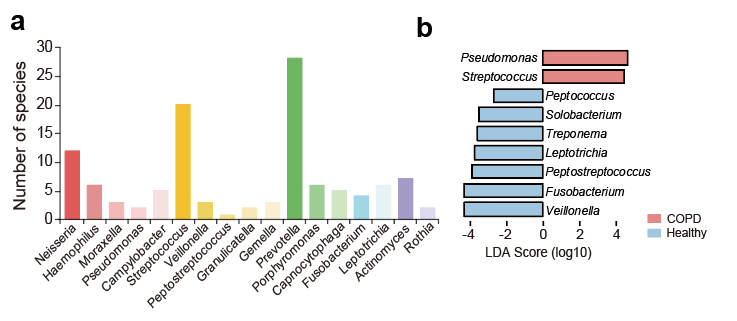

**Figure S2. a)** The number of species identified for each genus in the full-length 16S rRNA gene sequencing data. **b)** The 9 discriminatory genus-level taxa between COPD and controls identified using LEfSe (LDA>2.0).

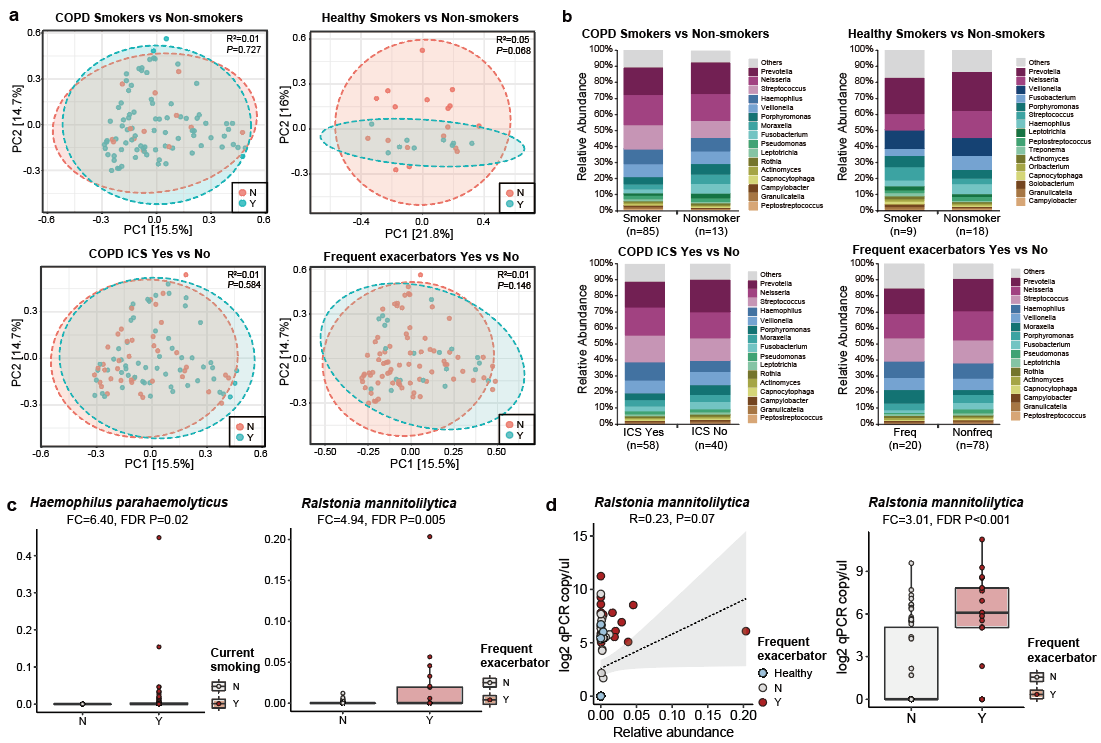

**Figure S3. a)** Principal coordinate analysis plots and **b)** genus-level microbiome profiles between COPD smokers versus non-smokers, healthy smokers versus non-smokers, COPD inhaled corticosteroid takers versus non-takers, and COPD frequent (defined as exacerbation events >=2/last year) and non-frequent exacerbators. **c)** Significant increase of *Haemophilus parahaemolyticus* in COPD smokers verus non-smokers (Fold-change=6.40, FDR *P*=0.02). Significant increase of *Ralstonia mannitolilytica* in frequent versus non-frequent exacerbators (Fold-change=4.94, FDR *P*=0.005, Figure S3c). **d)** qPCR assay based on species-specific primers for *Ralstonia mannitolilytica* confirmed the sequencing results.

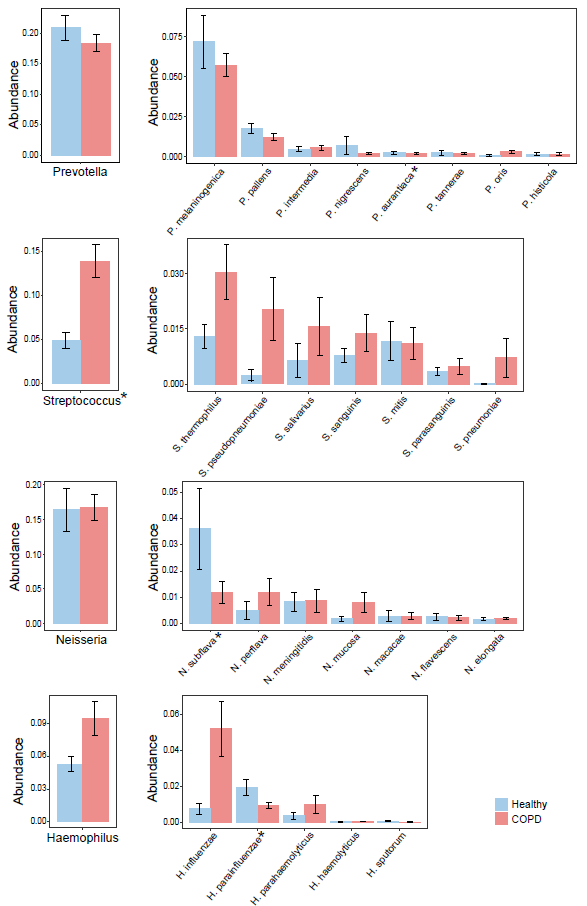

**Figure S4.** Heterogeneity in the changes in relative abundance of individual species (relative abundance>0.005) within *Prevotella*, *Streptococcus*, *Neisseria* and *Haemophilus* in COPD versus healthy controls. * FDR *P*<0.05

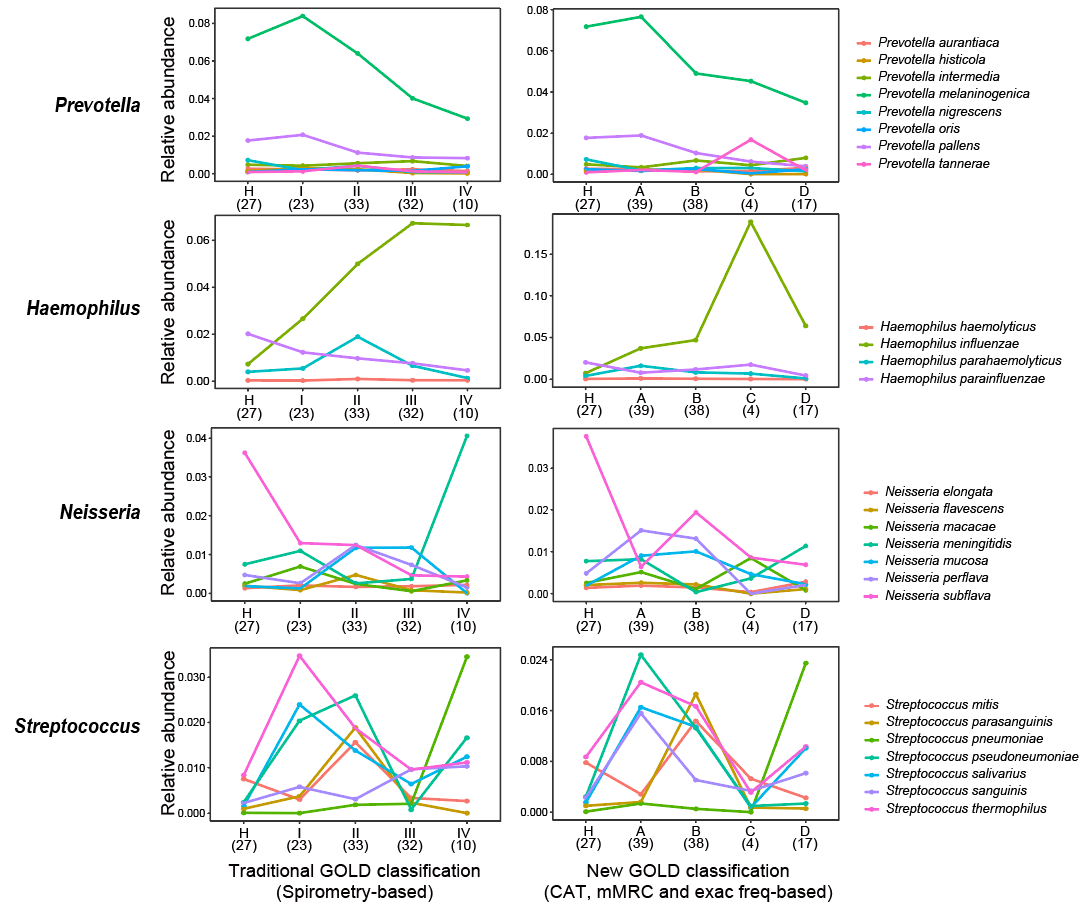

**Figure S5.** The alternation of individual species (relative abundance>0.005) within *Prevotella*, *Haemophilus*, *Neisseria* and *Streptococcus* in healthy controls and in COPD patients with increased disease severity. The COPD patients were classified based on traditional GOLD classification (spirometry-based) and the new GOLD classification scheme in the 2019 guideline (based on mMRC, CAT score and exacerbation frequency). The number of subjects in each subgroup was indicated in the parenthesis.

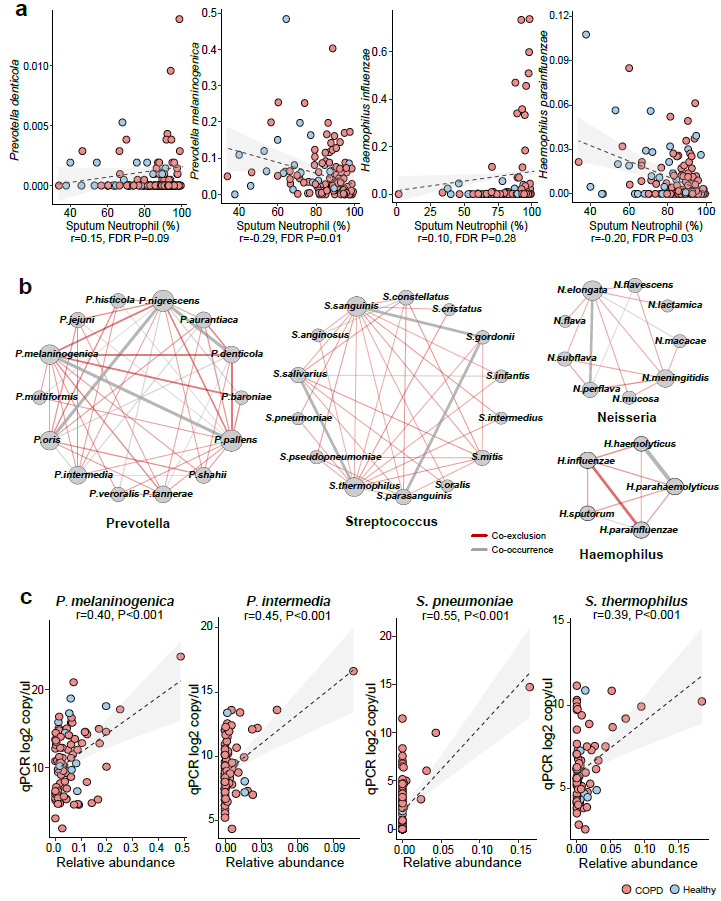

**Figure S6.** Intra-genus heterogeneity of sputum microbiome in COPD and healthy subjects. **a)** The reciprocal relationship between *Prevotella denticola* and *Prevotella melaninogenica*, and between *Haemophilus influenzae* and *Haemophilus parainfluenzae* and sputum neutrophilic percentage. **b)** More pervasive co-exclusive versus co-occurrence relationships between individual species within *Prevotella*, *Streptococcus*, *Neisseria* and *Haemophilus*. Only significant correlations were shown in the networks (SparCC, *P*<0.05). Co-exclusion relationships were in red, whereas co-occurrence relationships were in grey. **c)** Concordance between absolute copy number of *P. melaninogenica*, *P. intermedia*, *S. pneumoniae*, and *S. thermophilus* as revealed by qPCR assays and their corresponding relative abundances in full-length 16S sequencing.

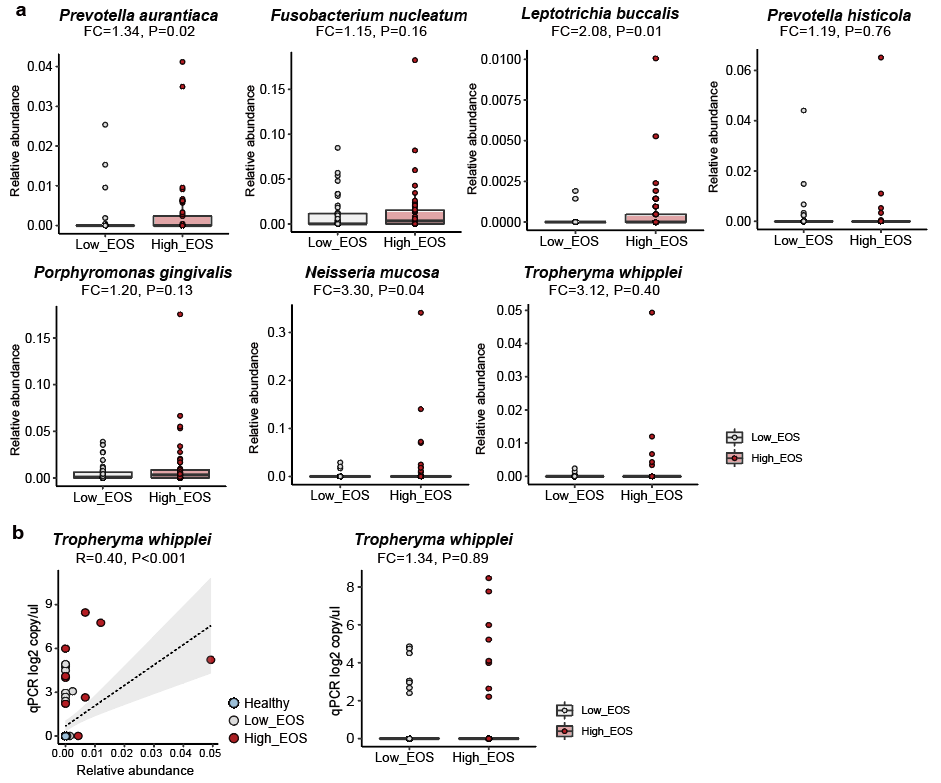

**Figure S7.** **a)** The alternations of the seven bacterial species associated with eosinophilic inflammations between patients with low eosinophilic levels (sputum EOS<3%) and high eosinophilic levels (sputum EOS>=3%). **b)** qPCR assays on *T. whipplei* confirmed the sequencing results.

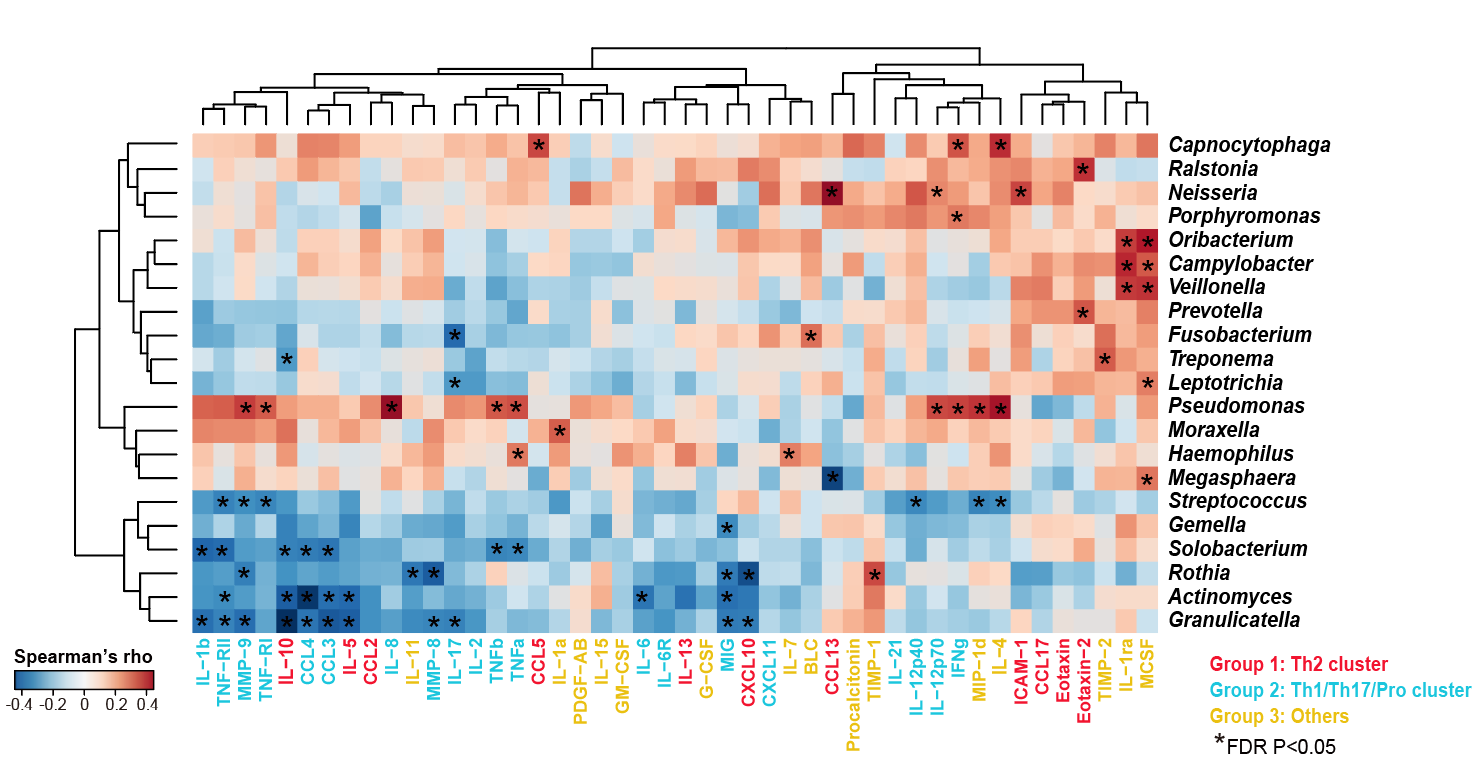

**Figure S8.** Clustered heatmap on correlations between genus-level microbiome profiles with the panel of 47 sputum mediators from a subset of 59 COPD patients. The genera were shown if they had relative abundance>0.001 and were significantly associated with at least one of the 47 sputum mediators (HAllA, FDR *P*<0.05). The significant correlations were indicated in asterisks. The 47 mediators were colored based on the assigned clusters (Group 1-3) from the correlation profile with the species-level microbiome features (Figure 3). No inflammatory phenotype-related clusters were observed for the genus-level associations.

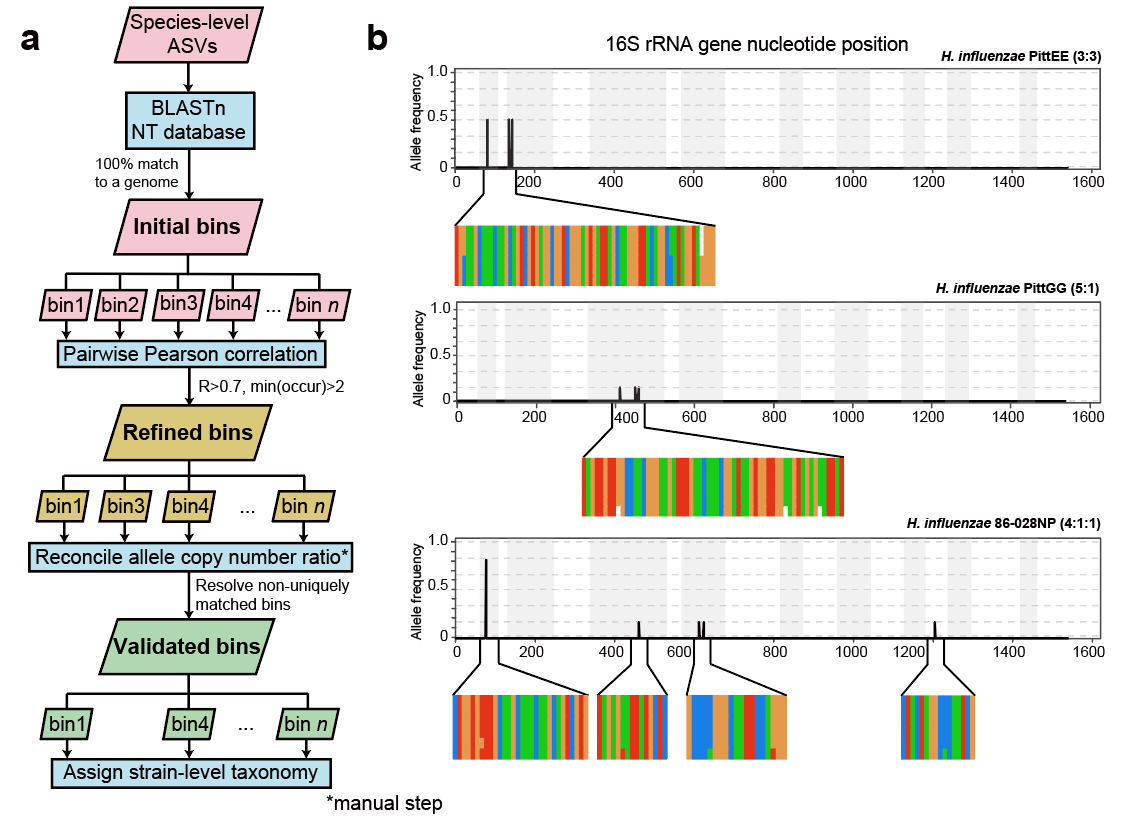

**Figure S9. a)** The designed semi-automated pipeline in identifying strain-level ASV bins. **b)** The polymorphisms in the 16S rRNA gene sequences for *H. influenzae* PittEE, PittGG and 86-028NP strains. The position and frequency of substitution in the full-length 16S sequences of the three strains were shown. Magnified regions showed respective positions in the alignment of all six copies of 16S gene in the corresponding *H. influenzae* genomes. The genuine ratios of 16S allelic variants in the three genomes are: 3:3, 5:1 and 4:1:1.

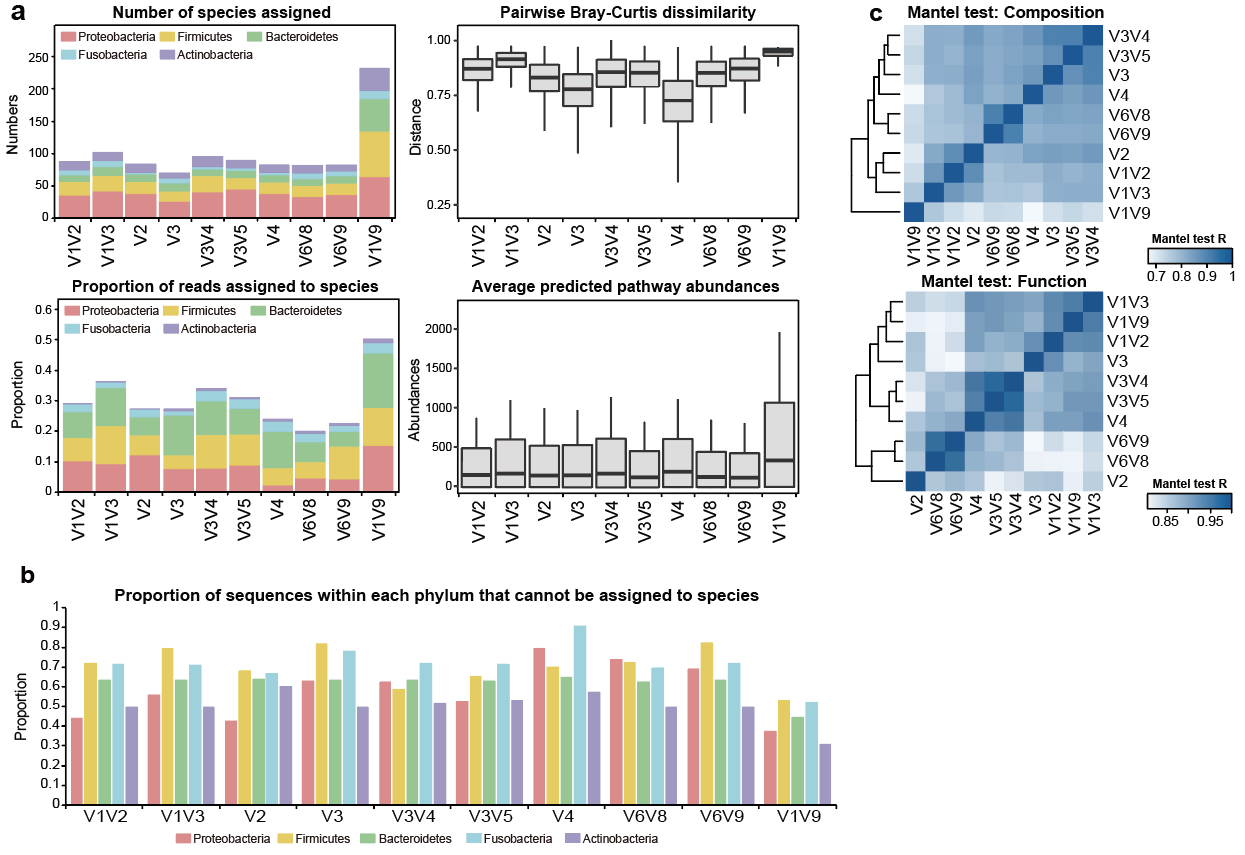

**Figure S10. a)** Comparisons between the full-length 16S data and the data for the partitioned 16S hypervariable regions, in terms of 1) the number of species-level taxa assigned, 2) the proportion of reads that can be assigned to species, 3) the pairwise Bray-Curtis dissimilarity across all samples, 4) the average abundances of pathways predicted by PICRUSt2. **b)** The proportion of sequences within each major phylum (Proteobacteria, Firmicutes, Bacteroidetes, Actinobacteria and Fusobacteria) that cannot be assigned with species-level taxa for the full-length sequences (V1V9) and nine individual hypervariable regions. **c)** Heatmap showing similarity in compositional and functional profiles between the full-length 16S data and data for individual 16S sub-regions. Mantel test was performed to assess the profile-level correlations.

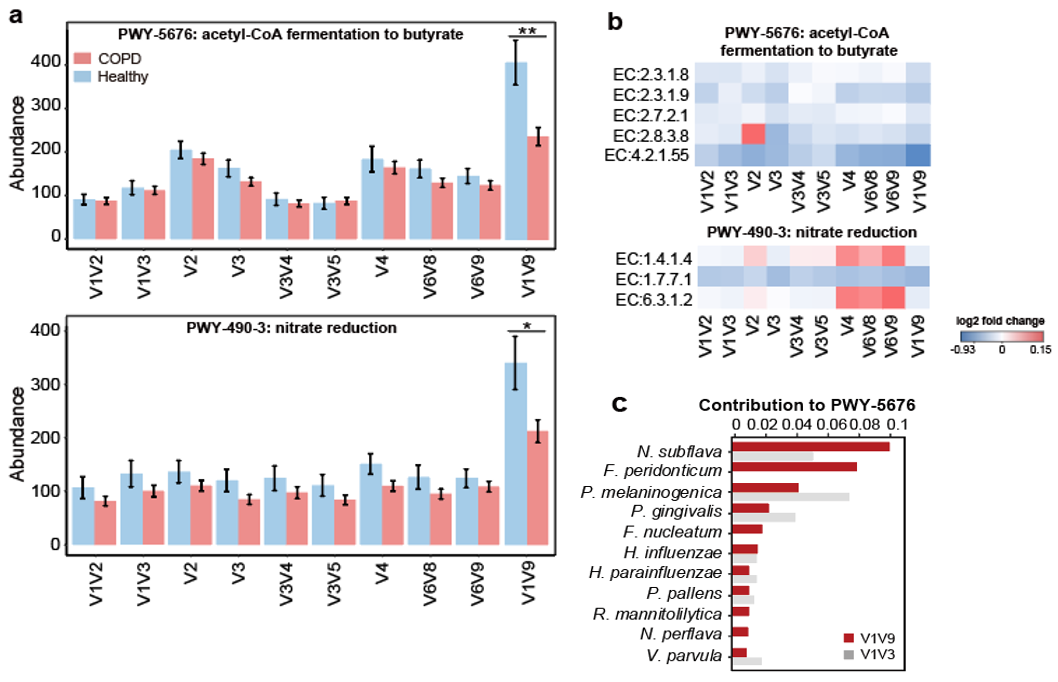

**Figure S11.** Functional inference using full-length 16S sequences and individual hypervariable regions using PICRUSt2. **a)** The inferred abundance in COPD and controls of pathways PWY-5676 (acetyl-CoA fermentation to butyrate) and PWY-490-3 (nitrate reduction) using the full-length 16S data (V1V9) and data for individual hypervariable regions. **b)** The inferred fold changes of individual genes within each pathway in COPD patients versus healthy controls. **c)** The top contributing bacterial species and their proportions of contribution to the ‘PWY-5676’ pathway as inferred from V1V9 and V1V3 data.

**Table S1.** The number of species-level taxa identified using the same pipeline (the DADA2 pipeline customized for 454 or Illumina platforms) for all previous COPD airway microbiome datasets sequencing individual hypervariable regions of the 16S rRNA gene.

| **Accession** | **Sample type** | **PMID** | **Platform** | **16S region** | **Sample size** | **Number of species identified** |
| --- | --- | --- | --- | --- | --- | --- |
| SRP102480 | Sputum | 29269441 | Miseq | V4 | 446 | 104 |
| SRP102629 | Sputum | 29386298 | Miseq | V4 | 423 | 106 |
| SRP073159 | Sputum | 29101284 | Miseq | V3-V4 | 134 | 82 |
| SRP065072 | Sputum | 26917613 | 454 | V3-V5 | 106 | 116 |
| ERP108788 | Sputum | NA | Miseq | V3-V4 | 95 | 125 |
| Dyrad.5GC82 | Sputum | 28851370 | Miseq | V3-V4 | 81 | 120 |
| SRP066375 | Sputum | NA | Miseq | V4 | 181 | 112 |
| ERP022665 | BAL | 29992131 | 454 | V3-V5 | 64 | 82 |
| ERP003401 | Sputum | 25253795 | 454 | V1-V2 | 31 | 21 |
| ERP014054 | Sputum | NA | 454 | V6-V8 | 28 | 30 |
| SRP124904 | Sputum | 29518088 | 454 | V3-V4 | 26 | 18 |
| SRP064237 | Bronchial | 28704452 | 454 | V6-V9 | 39 | 41 |
| SRP065328 | BAL | 27486204 | Miseq | V4 | 20 | 13 |
| SRP057611 | Bronchial | 27146202 | Miseq | V1-V2 | 37 | 81 |
| SRP122946 | Bronchial | 29316977 | Miseq | V3 | 18 | 107 |
| SRP107187 | Sputum | 29579057 | 454 | V6-V8 | 14 | 54 |
| SRP075523 | Sputum | 27428540 | Miseq | V4 | 8 | 28 |
| SRP068430 | Bronchial | NA | Miseq | V2 | 12 | 28 |

**Table S2.** The list of strain-level ASV bins corresponding to 10 strain-level taxa identified using the pipeline designed in this study.

| **ASV bin ID** | **Species** | **Genome (Strain) best hit** | **ASVs in the bin** | **Pearson’s R** | **Genuine allele variants** |
| --- | --- | --- | --- | --- | --- |
| 363 | *Fusobacterium nucleatum* | subsp. vincentii | ASV1183;ASV835 | 0.97 | 1:1:1 |
| 396 | *Fusobacterium periodonticum* | KCOM 1261 | ASV141;ASV39 | 0.85 | 1:1:1 |
| 449 | *Haemophilus influenzae* | PittEE | ASV472;ASV1085 | 0.99 | 3:3 |
| 461 | *Haemophilus influenzae* | 86-028NP | ASV10;ASV36 | 0.97 | 4:1:1 |
| 490 | *Haemophilus influenzae* | PittGG | ASV182;ASV1449 | 0.93 | 5:1 |
| 534 | *Haemophilus parainfluenzae* | T3T1 | ASV272;ASV335 | 0.81 | 4:1:1 |
| 822 | *Prevotella jejuni* | CD3:33 | ASV180;ASV207 | 0.81 | 1:1:1:1 |
| 2074 | *Streptococcus pseudopneumoniae* | IS7493 | ASV30 | NA | 1 |
| 2171 | *Veillonella dispar* | NCTC11831 | ASV123;ASV191 | 0.72 | 1:1:1:1 |
| 2087 | *Streptococcus salivarius* | NCTC7366 | ASV17;ASV53 | 0.75 | 5:1 |

**Table S3.** The list of 9 PICRUSt-predicted pathways uniquely identified as significantly different in abundance between COPD and healthy controls by the full-length 16S data. For each pathway, their abundances, fold-changes and *P*-values were shown for the full-length (V1V9) data and data for the nine 16S hypervariable regions as average values. All these pathways were statistically significant in V1V9 data but non-significant in all nine sub-regions data.

| **Pathway ID** | **V1V9_abundance** | **Average_abundance** | **V1V9_FC** | **Average_FC** | **V1V9_FDR** | **Average FDR** | **Description** |
| --- | --- | --- | --- | --- | --- | --- | --- |
| PWY-5676 | 289.872 | 131.514 | -0.661 | -0.174 | 0.004 | 0.266 | acetyl-CoA fermentation to butyrate |
| PWY-5484 | 1347.833 | 718.903 | -0.399 | -0.121 | 0.007 | 0.304 | glycolysis II (from fructose 6-phosphate) |
| PWY490-3 | 256.695 | 107.023 | -0.544 | -0.112 | 0.028 | 0.219 | nitrate reduction |
| PWY-7219 | 1938.478 | 1006.231 | -0.276 | -0.106 | 0.030 | 0.396 | adenosine ribonucleotides de novo biosynthesis |
| GLYOXYLATE-BYPASS | 129.744 | 100.389 | 1.042 | 0.524 | 0.032 | 0.238 | glyoxylate cycle |
| THRESYN-PWY | 1550.343 | 800.959 | -0.260 | -0.096 | 0.035 | 0.446 | superpathway of L-threonine biosynthesis |
| PWY-6386 | 1741.622 | 897.568 | -0.274 | -0.105 | 0.037 | 0.397 | UDP-N-acetylmuramoyl-pentapeptide biosynthesis II |
| PWY-5695 | 1452.145 | 785.254 | -0.259 | -0.116 | 0.038 | 0.292 | urate biosynthesis/inosine 5'-phosphate degradation |
| PWY-6123 | 1790.831 | 922.248 | -0.183 | -0.114 | 0.041 | 0.320 | inosine-5'-phosphate biosynthesis I |

**Table S4.** The average and standard deviation of Ct values for butyryl-CoA:acetate CoA-transferase gene (EC:2.8.3.8) and internal control (*rpoB* gene) in a subset of 88 COPD and healthy subjects. The values for ΔCt, ΔΔCt, 2^-ΔΔCt (COPD vs Healthy) and decrease fold change are shown.

|  | **EC:2.8.3.8** | **rpoB gene** |
| --- | --- | --- |
| COPD (Ct) | 33.17±3.36 | 25.97±3.03 |
| Healthy (Ct) | 31.18±3.43 | 26.09±3.05 |
| COPD (ΔCt) | 8.20±3.00 |  |
| Healthy (ΔCt) | 6.09±2.24 |  |
| ΔΔCt (COPD vs Healthy) | 2.105 |  |
| 2^-ΔΔCt | 0.232 |  |
| Decrease folds | 4.317 |  |

**Table S5.** All the ASVs and their numbers of sequences identified in the reagent controls.

| **ASV** | **DNA extraction blank** | **PCR blank** | **Taxonomy** |
| --- | --- | --- | --- |
| ASV1 | 32 | 0 | k__Bacteria; p__Proteobacteria; c__Alphaproteobacteria; o__Rhizobiales; f__Hyphomicrobiaceae; g__Devosia; s__ |
| ASV2 | 12 | 18 | k__Bacteria; p__Proteobacteria; c__Alphaproteobacteria; o__Rhodobacterales; f__Rhodobacteraceae; g__; s__ |
| ASV3 | 8 | 16 | k__Bacteria; p__Planctomycetes; c__Phycisphaerae; o__MSBL9; f__; g__; s__ |
| ASV4 | 5 | 9 | k__Bacteria; p__Proteobacteria; c__Gammaproteobacteria; o__Thiotrichales; f__Piscirickettsiaceae; g__; s__ |
| ASV5 | 2 | 1 | k__Bacteria; p__Proteobacteria; c__Deltaproteobacteria; o__Syntrophobacterales; f__Syntrophobacteraceae; g__; s__ |
| ASV6 | 1 | 0 | k__Bacteria; p__Chloroflexi; c__S085; o__; f__; g__; s__ |
| ASV7 | 0 | 2 | k__Bacteria; p__OP11; c__WCHB1-64; o__; f__; g__; s__ |

**Table S6.** Occurrence and average relative abundance of contaminate genera detected in sequenced negative ‘blank’ controls by Salter et al.[6] in our microbiome data. The first column (occurrence at relative abundance>0) was calculated as the fraction of samples in which each genus has abundance greater than 0. The second column (occurrence at relative abundance>0.1) was calculated as the fraction of samples in which each genus has abundance greater than 0.1. And the third column is the average relative abundance of each genus across all samples.

| **Genus** | **Occurrence (at relative abundance > 0)** | **Occurrence (at relative abundance > 0·1)** | **Average relative abundance** |
| --- | --- | --- | --- |
| Alphaproteobacteria |  |  |  |
| *Afipia* | 0.008 | 0 | 3.83E-06 |
| *Aquabacterium* | 0.006 | 0 | 2.61E-05 |
| *Asticcacaulis* | 0 | 0 | 0 |
| *Aurantimonas* | 0 | 0 | 0 |
| *Beijerinckia* | 0 | 0 | 0 |
| *Bosea* | 0 | 0 | 0 |
| *Bradyhizobium* | 0 | 0 | 0 |
| *Brevundimonas* | 0 | 0 | 0 |
| *Caulobacter* | 0 | 0 | 0 |
| *Craurococcus* | 0 | 0 | 0 |
| *Devosia* | 0.096 | 0 | 0.000137931 |
| *Hoeflea* | 0 | 0 | 0 |
| *Mesorhizobium* | 0 | 0 | 0 |
| *Methylobacterium* | 0.024 | 0 | 1.15E-05 |
| *Novosphingobioum* | 0 | 0 | 0 |
| *Ochrobactrum* | 0.008 | 0 | 0.00027969 |
| *Paracoccus* | 0.006 | 0 | 1.53E-05 |
| *Pedomicrobiom* | 0 | 0 | 0 |
| *Phyllobacterium* | 0.008 | 0 | 0.000180077 |
| *Rhizobium* | 0 | 0 | 0 |
| *Roseomonas* | 0 | 0 | 0 |
| *Sphingobium* | 0 | 0 | 0 |
| *Sphingomonas* | 0.006 | 0 | 0.000229885 |
| *Sphingopyxis* | 0 | 0 | 0 |
| Betaproteobacteria |  |  |  |
| *Acidovorax* | 0 | 0 | 0 |
| *Azoarcus* | 0 | 0 | 0 |
| *Azospira* | 0 | 0 | 0 |
| *Burkholderia* | 0 | 0 | 0 |
| *Comamonas* | 0 | 0 | 0 |
| *Cupriavidus* | 0 | 0 | 0 |
| *Curvibacter* | 0 | 0 | 0 |
| *Delftia* | 0 | 0 | 0 |
| *Duganella* | 0 | 0 | 0 |
| *Herbaspirillum* | 0 | 0 | 0 |
| *Janthinobacterium* | 0 | 0 | 0 |
| *Kingella* | 0 | 0 | 0 |
| *Leptothrix* | 0 | 0 | 0 |
| *Limnobacter* | 0 | 0 | 0 |
| *Massilia* | 0.008 | 0 | 7.66E-06 |
| *Methylophilus* | 0 | 0 | 0 |
| *Methyloversatilis* | 0 | 0 | 0 |
| *Oxalobacter* | 0 | 0 | 0 |
| *Pelomonas* | 0.032 | 0 | 3.83E-05 |
| *Polaromonas* | 0 | 0 | 0 |
| *Ralstonia* | 0.416 | 0.008 | 0.005724 |
| *Schlegelella* | 0 | 0 | 0 |
| *Sulfuritalea* | 0 | 0 | 0 |
| *Undibacterium* | 0 | 0 | 0 |
| *Variovorax* | 0 | 0 | 0 |
| Gammaproteobacteria |  |  |  |
| *Acinetobacter* | 0.416 | 0 | 0.001916 |
| *Enhydrobacter* | 0.008 | 0 | 7.66E-06 |
| *Enterobacter* | 0 | 0 | 0 |
| *Escherichia* | 0 | 0 | 0 |
| *Nevskia* | 0 | 0 | 0 |
| *Pseudomonas* | 0.28 | 0.016 | 0.016146 |
| *Pseudoxanthomonas* | 0 | 0 | 0 |
| *Psychobacter* | 0 | 0 | 0 |
| *Stenotrophomonas* | 0.032 | 0 | 1.53E-05 |
| *Xanthomonas* | 0 | 0 | 0 |
| Actinobacteria |  |  |  |
| *Aeromicrobium* | 0 | 0 | 0 |
| *Arthrobacter* | 0 | 0 | 0 |
| *Beutenbergia* | 0 | 0 | 0 |
| *Brevibacterium* | 0 | 0 | 0 |
| *Corynebacterium* | 0.004 | 0 | 0.000843 |
| *Curtobacterium* | 0 | 0 | 0 |
| *Dietzia* | 0.008 | 0 | 3.83E-06 |
| *Geodermatophilus* | 0 | 0 | 0 |
| *Janibacter* | 0 | 0 | 0 |
| *Kocuria* | 0 | 0 | 0 |
| *Microbacterium* | 0.006 | 0 | 0.00067 |
| *Micrococcus* | 0.006 | 0 | 7.66E-06 |
| *Microlunatus* | 0 | 0 | 0 |
| *Patulibacter* | 0 | 0 | 0 |
| *Propionibacterum* | 0 | 0 | 0 |
| *Rhodococcus* | 0.024 | 0 | 1.53E-05 |
| *Tsukamurella* | 0 | 0 | 0 |
| Firmicutes |  |  |  |
| *Abiotrophia* | 0.006 | 0 | 0.000153 |
| *Bacillus* | 0.032 | 0 | 3.07E-05 |
| *Brevibacillus* | 0 | 0 | 0 |
| *Brochothrix* | 0 | 0 | 0 |
| *Facklamia* | 0 | 0 | 0 |
| *Paenibacillus* | 0 | 0 | 0 |
| *Streptococcus* | 0.896 | 0.368 | 0.119103 |
| Bacteroidetes |  |  |  |
| *Chryseobacterium* | 0.032 | 0 | 4.98E-05 |
| *Dyadobacter* | 0 | 0 | 0 |
| *Flavobacterium* | 0 | 0 | 0 |
| *Hydrotalea* | 0 | 0 | 0 |
| *Niatella* | 0 | 0 | 0 |
| *Olivibacter* | 0 | 0 | 0 |
| *Pedobacter* | 0 | 0 | 0 |
| *Wautersiella* | 0 | 0 | 0 |
| Deinococcus |  |  |  |
| *Deinococcus* | 0 | 0 | 0 |

**Table S7.** The list of species-specific and strain-specific primers used in this study.

| **Species and strain** | **Target gene** | **Forward primer** | **Reverse primer** | **Size (bp)** |
| --- | --- | --- | --- | --- |
| *H. influenzae* | WP_005652235 outer membrane protein P6 | TTGGCGGWTACTCTGTTGCT | TGCAGGTTTTTCTTCACCGT | 296 |
| *H. parainfluenzae* | WP_014063976 hypothetical protein | TTCTACAGGCGGCCAAACGG | TCGGTTTCTCATCGGGTGGCA | 114 |
| *S. pneumoniae* | WP_001284361 pneumolysin | AGCGATAGCTTTCTCCAAGTGG | CTTAGCCAACAAATCGTTTACCG | 75 |
| *S. thermophilus* | WP_011225905 agmatinase | AGAATTCCAGGAGCGAGATTTGC | CCTCTCGAACAGTGGCCTTTGC | 223 |
| *P. melaninogenica* | WP_013264642 lipoprotein | TCGCTATTGGTGGTGAGGCTGA | TGCTTTTGTCGTACTGGTTGCGC | 104 |
| *P. intermedia* | WP_014708742 hypothetical protein | GCTCACAGAGGGTAGCGTGC | AGCTGCCCCAAATTGCCACC | 101 |
| *H. influenzae* PittEE | WP_005686347 hypothetical protein | TCAGTACTTTCGGCAACGTGGT | AGAGGCTACAGGAATCGGAGGA | 175 |
| *H. influenzae* PittGG | WP_012054916 AlpA family phage regulatory protein | AGCGAGGCAATTTTCCGAAGC | GCCATGCTGCGCCTCTTGTT | 119 |
| *H. influenzae* 86-028NP | WP_005672606 hypothetical protein | GCCTTACTGCCGTTTGTTTCGCA | GCACCGTCAGCTCCCTATGCA | 115 |
| *Ralstonia mannitolilytica* | AJW44311.1 hypothetical protein | ATGCCCCATTCCGTCAGCTT | AACACGCGCATCCCATGAAG | 154 |
| *T. whipplei* | WP_011102574 heat shock protein 65 | TGACGGGACCACAACATCTG | ACATCTTCAGCAATGATAAGAGAAGTT | 503 |
